# A behavior-manipulating virus relative as a source of adaptive genes for parasitoid wasps

**DOI:** 10.1101/342758

**Authors:** D. Di Giovanni, D. Lepetit, M. Boulesteix, Y. Couté, M. Ravallec, J. Varaldi

## Abstract

To circumvent host immune response, some hymenopteran endo-parasitoids produce virus-like structures in their reproductive apparatus that are injected into the host together with the eggs. These viral-like structures are absolutely necessary for the reproduction of these wasps. The viral evolutionary origin of these viral-like particles has been demonstrated in only a few cases of wasp species all belonging to the Ichneumonoidea superfamily. In addition, the nature of the initial virus-wasp association remains unknown for all. This is either because no closely related descendant infects the wasps, because it has not been sampled yet, or because the virus lineage went extinct. In this paper, we show that the virus-like particles (VLPs) produced by endoparasitoids of *Drosophila* belonging to the *Leptopilina* genus (superfamily Cynipoidea) do have a viral origin, solving the debate on their origin. Furthermore, the ancestral donor virus still has close relatives infecting one of the wasp species, thus giving us insights on the ecological interaction that possibly allowed the domestication process. Intriguingly, this contemporary virus is both vertically and horizontally transmitted and has the particularity to manipulate the superparasitism behavior of the wasp. This raises the possibility that behavior manipulation has been instrumental in the birth of such association between wasps and viruses.

## 1 Introduction

Genetic information is typically passed on from generation to generation through reproduction, *ie* vertical transmission. However, at some point during the course of evolution, organisms may gain DNA from unrelated organisms, through horizontal gene transfer (HGT). Most horizontally acquired DNA is probably purged from the genomes of the population either because it did not reach the germinal cells in case of metazoan species and/or because no advantage is carried by the foreign sequence. However, natural selection may retain the foreign DNA leading ultimately to genetic innovation in the population/species [38].

The high frequency and relevance of such phenomenon has been recognized for decades for bacteria but was considered to have had a marginal impact on the evolution of metazoans[40]. However, this view has been recently challenged due to the discovery of numerous examples of HGT in metazoans with some of them leading to genetic innovation[8]. The most notorious example involves retroviral envelope genes that have been endogenized, domesticated and multiply replaced in mammalian genomes[45]. In this case, the fusogenic and immunosuppressive properties of these viral proteins (syncitins) have been repeatedly recruited to permit the evolution of placental structures during mammalian diversification. Interestingly, a similar case of syncitin domestication was recently described in a clade of viviparous Scincidae lizards that also rely on a placenta-like structure to feed their offspring [20]. Other examples include phytophagous mites and Lepidoptera that deal with chemical defenses of their host plant thanks to the acquisition of a bacterial gene involved in detoxification [79], several phytophagous arthropods (Aphids, mites and gall midges) who independently acquired genes involved in carotenoid biosynthesis from fungal donors[58][31][15], or parasitic nematodes that domesticated plant cell-wall degrading enzymes from bacteria[21].

Regarding the question of domestication of horizontally-transfered DNA in eukaryotes, endoparasitic wasps are of particular interest because they have repeatedly domesticated not only single genes but entire viral machineries (review in [27] and since then [10]). Endoparasitic wasps lay their eggs inside the body of other arthropods, usually other insects, ultimately killing them. Their progeny is thus exposed to the host immune system. Notably, it has been found that the ancestor of at least three monophyletic groups of endoparasitic wasps have independently domesticated a battery of viral genes allowing them to deliver either DNA encoding immuno-suppressive factors or immuno-suppressive proteins themselves [4][78][10]. Strikingly, in the case DNA is delivered into the host (so-called polydnaviruses, PDV), it integrates into the host hemocytes DNA and gets expressed [5][14], manipulating the host physiology and behavior, ultimately favoring the development of wasp offspring. In cases where proteins are delivered, the viral machinery permits the delivery of these virulence proteins into host immune cells, thus inhibiting the host immune response[67]. In both cases, virally-derived genes are used by the wasp to produce a vector toolset composed of capsids and/or envelopes. However, the virulence factors themselves (or the DNA encoding the virulence factors) are of eukaryotic origin, probably pre-dating the domestication event. Evolution has thus repeatedly favored the domestication of kits of viral genes allowing the production of virus-like structures in the reproductive apparatus of parasitic wasps with clear functional convergence.

One striking pattern emerging from the data, is that all described cases documented so far involve wasps belonging to the Ichneumonoidea superfamily [27]. Although this super-family is very speciose (most likely around 100,000 species), it represents a modest fraction of parasitic Hymenoptera diversity (most likely around 1 million species[25]). Another feature of the current data is that the biology of the ancestral donor virus is completely unclear. For one such domestication event (in the Campopleginae sub-family, Ichneumonidae family), the ancestral virus has not been identified at all, whereas a beta nudivirus has been identified as the donor virus for wasps belonging to the microgastroid complex of the Braconidae family[4]. In *Venturia canescens* (Campopleginae sub-family, Ichneumonidae family), the unique case of viral replacement documented so far, and in some wasp species from the genus *Fopius* (subfamily Opiinae, Braconidae family), it has been shown that an alpha-nudivirus was the donor[65][10]. However, close relatives of the donor viruses are not known to infect present-day wasps, nor to infect their hosts. One possible explanation is that the “donor” viral lineages went extinct and/or have not been sampled yet. The exact nature of the association wasp/virus that permitted such massive domestication events is thus still unclear.

In this work, we identify a new independent case of virus domestication in the genus *Leptopilina* which belongs to a very distantly related wasp superfamily (Cynipoidea, Figitidae) compared to all previously described cases. Those wasps are parasitoids of *Drosophila* larvae. We show that the genes of viral origin permit all *Leptopilina* wasp species to produce so-called virus-like particles (VLPs). VLPs have been known for decades in this genus[68]. They are produced in the venom gland of the wasp, are devoid of DNA but contain virulence proteins that are injected, together with the egg, into the *Drosophila* larva[19]. They protect wasp eggs from *Drosophila* immune response [68][18]. We show that a close relative of the ancestral donor virus is still segregating in the species *L. boulardi* and its biology has been extensively studied by our group[53][62][54][48][76]. The virus, known as LbFV, belongs to a possibly new dsDNA virus family related to Hytrosaviridae, and more distantly related to Nudiviridae and Baculoviridae[48]. The virus is vertically transmitted and manipulates the wasp behaviour by forcing infected females to lay their eggs into already parasitized larvae. This virus-induced “host-sharing” benefits to the virus since it allows its horizontal transmission to new parasitoid lineages. On the contrary, this “superparasitism” behaviour comes with a cost to wasp fitness, making it a nice example of behaviour manipulation[26]. This result suggests that heritable viruses such as LbFV, might have been instrumental in the birth of such association between wasps and viruses. In addition, it shows that virus domestication by parasitic wasps is not restricted to the Ichneumonoidea superfamily but may concern more diversity than previously thought.

## 2 Results

We analyzed the genomic sequences of *L. boulardi* [76], *L. clavipes* [41], *L. heterotoma* (this study) and a related species in the *Ganaspis* genus (*G. brasiliensis*, this study). All *Leptopilina* species as well as *G. brasiliensis* belong to the Figitidae family and are endoparasitoids developing from various species of *Drosophila*.

The basic statistics for the assemblies used in this paper are presented in table S1. With an N50 of 2080 bp the *G. brasiliensis* assembly appeared more fragmented than those from the *Leptopilina* species whose N50 ranges from 12807 bp to 17657 bp. This reflects its two to three times larger genome size likely due to its higher content in repetitive sequences (44.92% vs. 24.02-28.82%). All four genomes were sequenced with coverage depth above 24 (between 24x and 85x), which is most likely sufficient to get the whole gene set[51]. Accordingly, a BUSCO[71] analysis revealed that the vast majority of the 1066 single copy genes expected to be found in most arthropods are indeed present in all four assemblies (from 96.6% in *G. brasiliensis* to 99.1% in *L. boulardi*), making these assemblies suitable for HGT detection (table S1).

We inferred the relationships among the wasps under study using a set of 627 genes ubiquitous to all arthropods (see methods). As expected, the three *Leptopilina* species form a monophyletic clade with *L. heterotoma* being more closely related to *L. clavipes* than to *L.boulardi* (Fig. 1).

**Figure 1:**
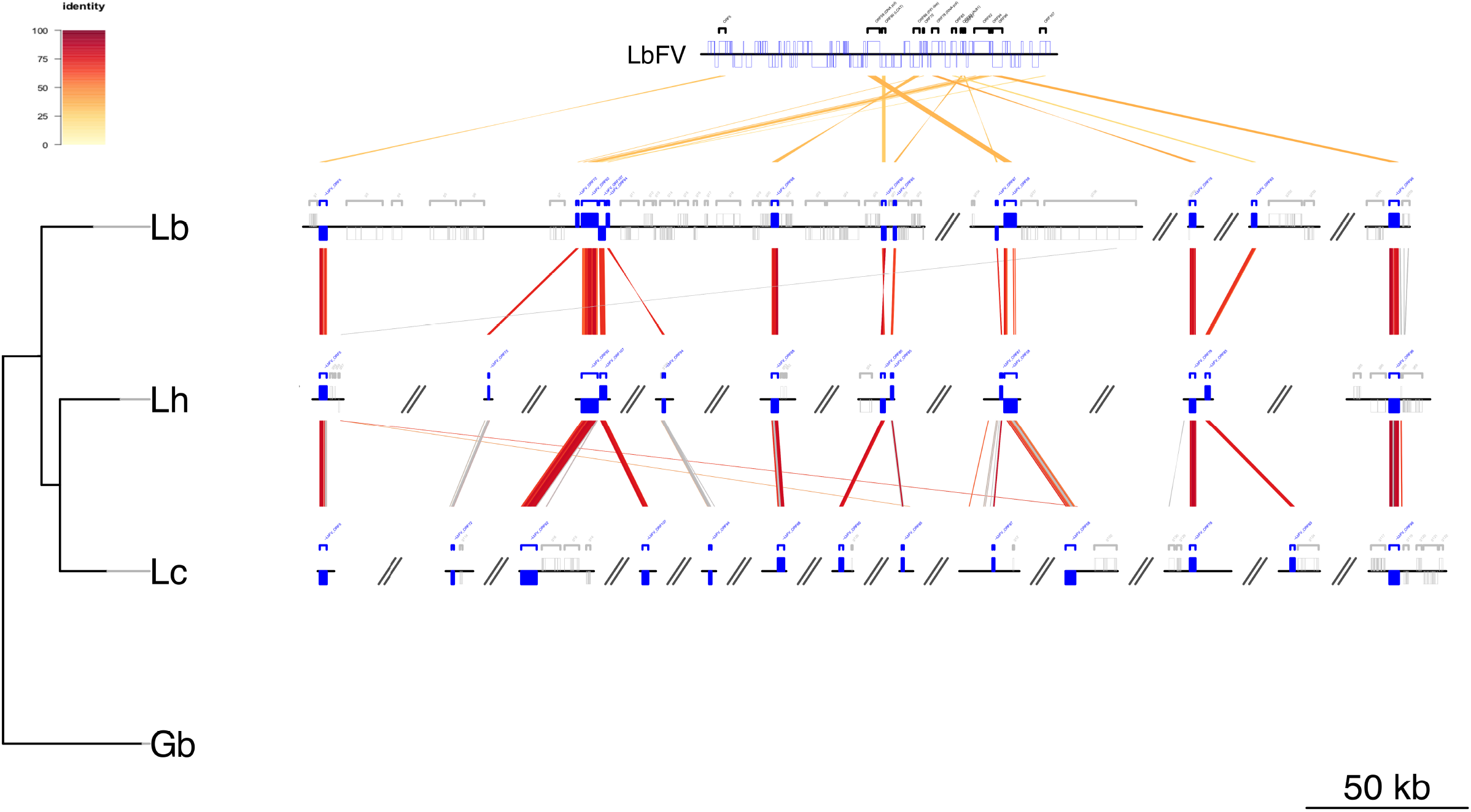
Comparative genomics of wasp scaffolds sharing similarities with virus proteins. Lb: *L. boulardi*, Lh: *L. heterotoma*, Lc: *L. clavipes*, Gb: *Ganaspis brasiliensis*, LbFV: Leptopilina boulardi Filamentous Virus. The species-tree on the left has been obtained using a concatenation of 627 universal arthropod genes. All branches (Lh-Lc and Lh-Lc/Lb) have an aLRT value of 1 (*Apis mellifera* was used as an outgroup). The red/yellow color code depicts the percentage of protein identity between homologous sequence pairs (viral or virally-derived loci). Grey connections indicate homology between regions that does not contain virally-derived loci. Genes of eukaryotic origin are depicted in grey on the scaffolds. The figure has been drawn using the genoPlotR package[34]. The scaffolds are ordered from left to right in an arbitrary manner.

In order to identify putative horizontal transfers between an LbFV-like virus and the wasps, we blasted the 108 proteins encoded by the behaviour-manipulating virus that infects *L. boulardi* (LbFV) against the *Leptopilina* and *Ganaspis* genomes (tblastn). Interestingly, we found that 17 viral proteins had highly significant hits in wasp genomes (1.3 × 10^−178^ < e-values < 10^−5^). Among them, two classes should be distinguished. The first class is composed of four viral genes (ORFs 11, 13, 27 and 66) that have strong similarities with both *Leptopilina* and *Ganaspis* genes (Fig. S1). We previously reported that these genes have probably been acquired horizontally by the virus from an ancestral insect before the *Leptopilina* diversification ([48], Fig. S1 & S2A). Two of them (ORFs 27 and 66) are predicted to encode inhibitors of apoptosis, whereas ORFs 11 and 13 encode a putative demethylase [48]. These two last genes may derive from a single horizontal transfer followed by a subsequent gene duplication [48]. In the following section, we will focus on the second class of genes identified by this blast analysis.

### 2.1 *Leptopilina* species captured 13 viral genes from an LbFV-like virus

More surprisingly, we found clear evidence that a single massive integration of viral DNA into wasp genomes occurred before the diversification of the *Leptopilina* genus and after the divergence between *Ganaspis* and *Leptopilina*. This event led to the integration of 13 viral genes into the genome of the wasps (Fig. S2B). The corresponding 13 viral proteins have highly significant hits with all *Leptopilina* species (4.10^−4^ < e-values < 1.310^−178^, median = 10^−33^), but not with *G. brasiliensis*. The percentages of identity between these 13 LbFV proteins and *Leptopilina* homologs ranged from 21.9 to 41.9 (table 1 and fig. S3-S15). All 13 loci displayed complete open reading frame (ORF) starting with a methionine and ending with a stop codon in the three wasp species, and their length was very similar to the corresponding ORF in LbFV genome (supplementary tables S2, S3 and S4; the regression slopes of ORF length in the wasp versus ORF length in LbFV were respectively 0.95, 1.02 and 0.894 for *L. boulardi, L. heterotoma* and *L. clavipes*; all *R*^2^ > 0.95 and all p-values< 10^−9^ on 11 d.f.). This suggests that those genes do not contain intron.

**Table 1:**
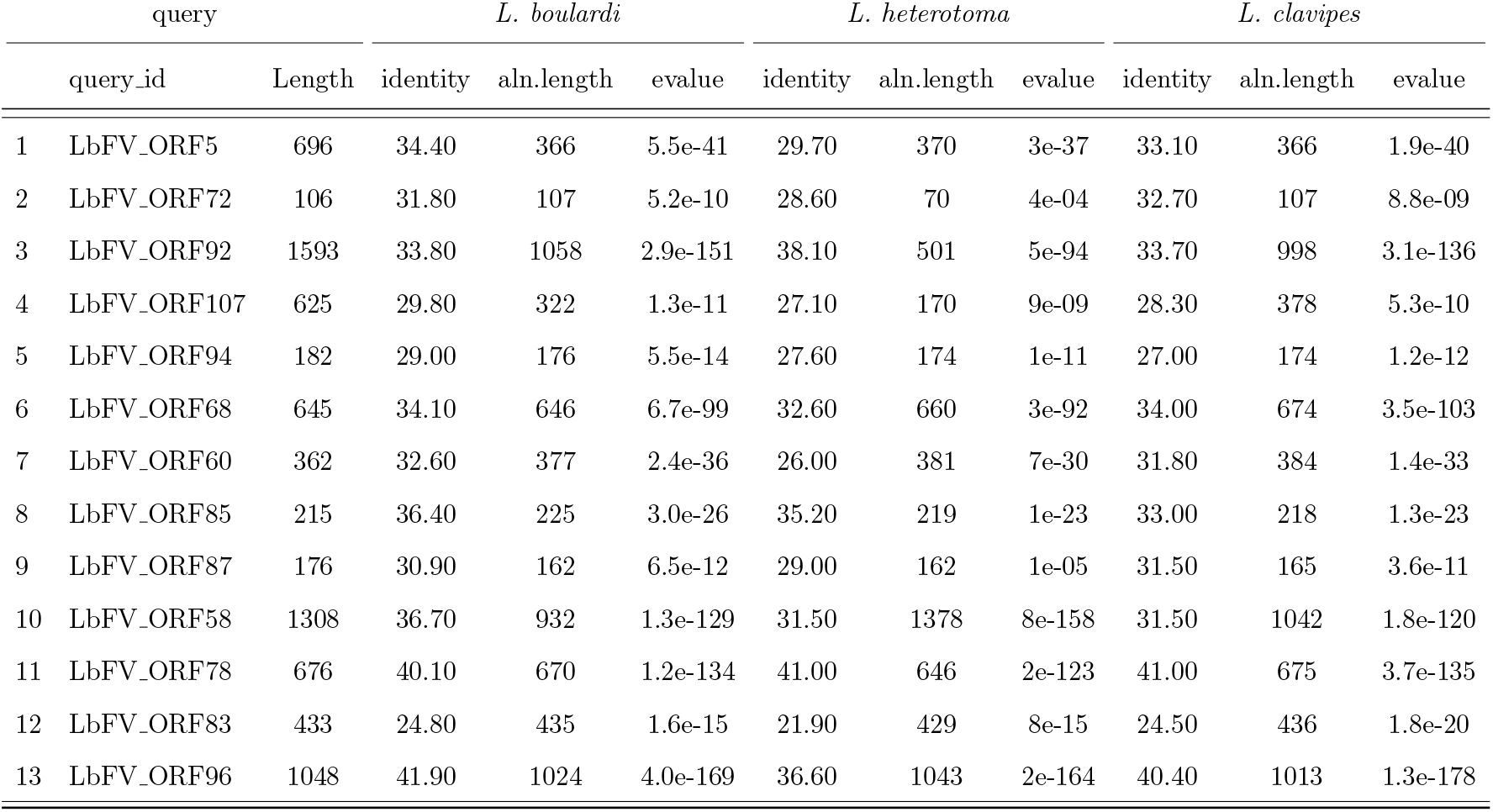
Blast hits for the 13 viral proteins against *Leptopilina* genomes (tblastn).

To define a set of expected features for typical scaffolds belonging to wasp genomes, we calculated the GC content and sequencing depth for scaffolds containing single-copy arthropod-universal BUSCO genes (Fig. S16). This is important since it allows one to distinguish genetic entities that may take part of the sample that have been sequenced. GC usually varies according to genomes, and coverage depth is directly related to the relative concentration of the DNA sequence under consideration. Except for one *L. clavipes* scaffold (scf7180005174277) encoding an homolog of ORF68, the general features (GC, sequencing depth) of wasp scaffolds sharing similarities with LbFV proteins were very similar to those calculated for the BUSCO-containing scaffolds (tables S2, S3, S4 and fig. S16). On the contrary, by analysing these statistics (GC and coverage), we could easily detect the presence of some known extra-chromosomal symbionts such as the virus LbFV in *L. boulardi* (Fig. S16A), or the bacteria *Wolbachia* in *L. heterotoma* (Fig. S16B). In addition, several typical intron-containing eukaryotic genes were predicted in the vicinity of these genes (depicted in grey in Fig. 1). Note that apart from these 13 loci specifically found in *Leptopilina* genomes, most flanking *Leptopilina* predicted proteins were also detected in the *G. brasiliensis* genome (66/72 for *L. boulardi*, 8/11 for *L. heterotoma* and 10/15 for *L.clavipes*) showing that the absence of homologs in *G. brasiliensis* genome was not the consequence of a less reliable assembly. Taken together, these observations demonstrate that the *Leptopilina* scaffolds containing viral-like genes are part of the wasp genomes. The special case of scf7180005174277 in *L. clavipes* assembly may be the consequence of recent duplications for this gene, possibly explaining its higher coverage depth.

The evolutionary history of the thirteen genes is consistent with an horizontal transfer from an ancestor of the virus LbFV (or a virus closely related to this ancestor) to *Leptopilina* species (Figure 2). Indeed, when other sequences with homology to the proteins of interest were available in public databases, the three wasp genomes always formed a highly supported monophyletic clade with LbFV as a sister group of *Leptopilina* sequences (ORFs 58, 78, 92, 60, 68, 85, 96). In addition, for the 6 remaining phylogenies (for which no homologs was available in public databases), the mid-point rooting method always led to similar topologies with LbFV as the sister group of *Leptopilina* sequences. Furthermore, the divergence LbFV-*Leptopilina* relative to the divergence among *Leptopilina* species was identical for both types of loci (Fig. S17), further suggesting that both loci have the same evolutionary history. Interestingly, it appeared from this analysis of ORF60, that before being transfered to *Leptopilina* wasps, the gene has probably been acquired by the donor virus from an ancestral bacteria (Figure 2).

**Figure 2:**
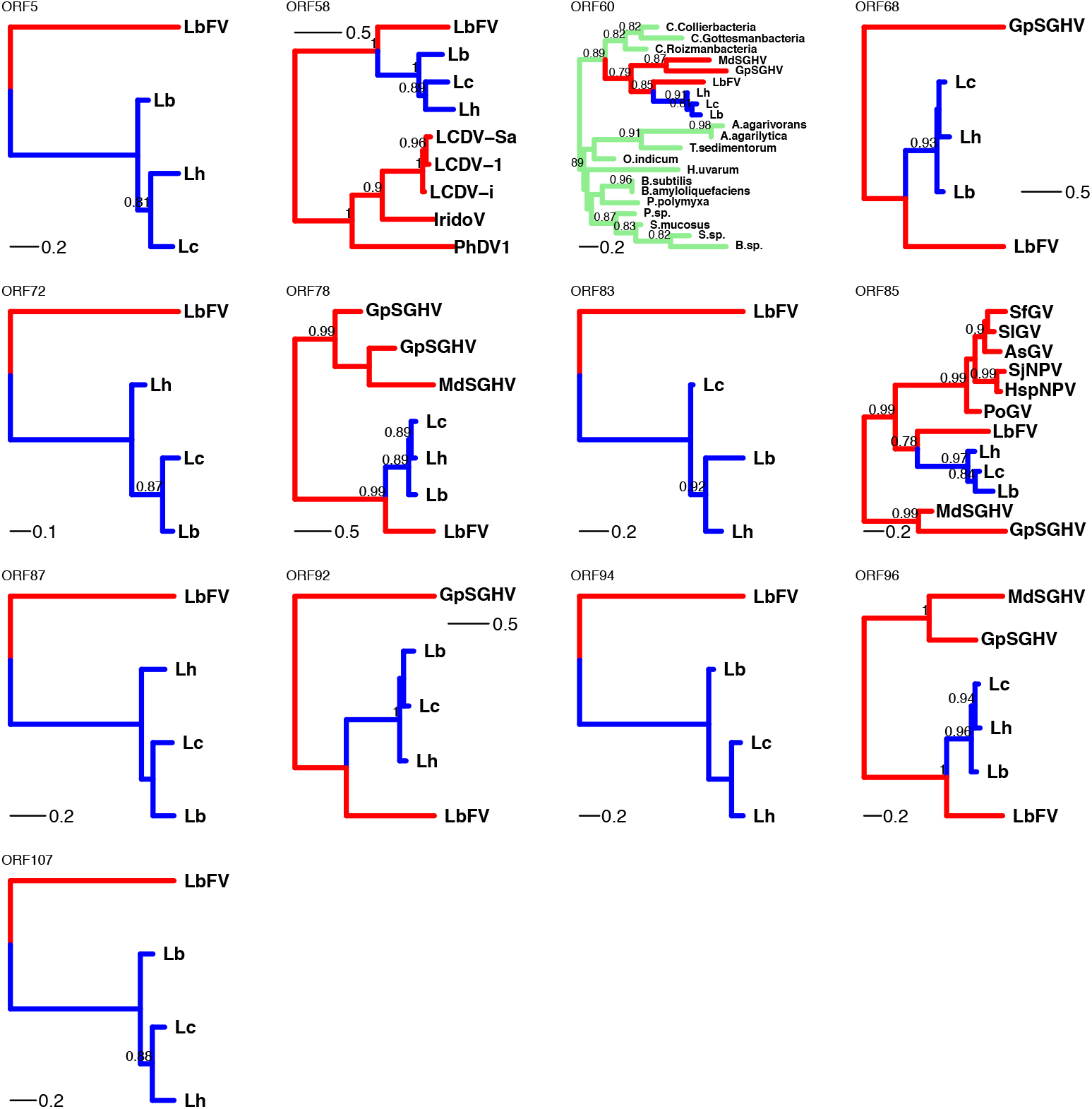
Phylogenetic evidence for a massive horizontal transfer of thirteen viral genes into the genome of *Leptopilina* wasps. The names of the ORFs refers to the ORF number in LbFV genome. Blue, red and green colors represent respectively (supposedly) eukaryotic, viral or bacterial branches. Only aLRT supports ⩾ 0.7 are shown. The mid-point rooting method was used. Accession numbers of the corresponding sequences are available in table S5.

The clustering of most of these loci on the same scaffold in *L. boulardi* (8 out of 13 on scaffold 159, N=75550 scaffolds, see Figure 1) strongly suggests that a single event is at the origin of the phenomenon. In addition, for a few pairs of *L. boulardi* and *L. heterotoma* scaffolds, it was possible to test for the synteny of their virally-derived genes (ORFs 92 and 107 in scaffolds 159 in Lb and IDBA_7081 in Lh, and ORFs 87 and 58 in scaffolds 2503 of Lb and IDBA_5653 in Lh). In all cases, the synteny appeared to be maintained between the two *Leptopilina* species (Fig. 1). In addition, a few flanking non-virally derived sequences were co-occuring around the same viral genes in different *Leptopilina* species (grey connections in Fig.1, see Fig. S18 for details). The overall shared organization of these genes in the three *Leptopilina* species suggests that they have been vertically inherited since a single ancestral endogenization event.

To further assess the distribution of those virally-derived genes in the diversity of *Leptopilina* wasps, we designed primers for ORF96 which is the most conserved gene. We successfully PCR amplified and sequenced the corresponding PCR product from DNA extracts obtained from all *Leptopilina* species tested (*L. guineaensis, L. freyae, L. victoriae* in addition to *L. boulardi, L. heterotoma* and *L. clavipes*, figure S19A). The phylogeny obtained after the sequencing of the PCR products was congruent with the species-tree estimated from a phylogeny based on ITS2 sequences (Fig. S19B). As expected, no PCR product was obtained from *Ganaspis brasiliensis* extracts.

### 2.2 Virally-derived genes are under strong purifying selection in wasp genomes

In order to assess the way natural selection have acted on these virally-derived genes since their endogenization, we calculated the dN/dS ratios using alignments involving the three *Leptopilina* species. We also calculated dNdS ratios for a set of 942 genes found in the three *Leptopilina* species and that are also shared by at least 90% of all arthropods[71]. Those genes are thus expected to be under strong purifying selection. Accordingly, the ‘universal” arthropod gene set had a very low dN/dS mean value (mean=0.114, median=0.085), with a distribution skewed towards 0 (Figure S20). Interestingly, the thirteen virally-derived genes had very low and very similar dN/dS values (mean=0.215, median=0.222, min=0.125, max=0.284), suggesting that they are all as essential for the survival and/or reproduction of *Leptopilina* wasps as any “universal” arthropod gene.

### 2.3 Virally-derived genes are only expressed in female venom glands at the onset of VLPs production

All *Leptopilina* species studied so far (*L. heterotoma, L. boulardi* and *L. victoriae*) produce VLPs in their venom gland [68][24][57]. As expected, we found that *L. clavipes* also produce VLPs in their venom gland, further suggesting that this is a general feature for all *Leptopilina* species (Fig. S21). Because VLPs are known to protect their eggs from *Drosophila* immune reaction in *Leptopilina* [68][42][57], we wondered whether the 13 virally-derived genes were in fact responsible for their production. Under this hypothesis, our prediction was that the 13 genes would be expressed only in the venom gland of females since VLPs are specifically produced in this tissue, and only when VLPs are being produced.

To test this idea, we measured the expression of the 13 virally-derived genes in the venom glands, ovaries, rest of the body of *L. boulardi* females, and also in *L. boulardi* males. We followed their expression from the very beginning of the pupal stage (day 11) until the emergence of the host (day 21). During that period, the venom gland is being formed and is matured (Fig. S22). The venom gland produces the VLPs that are released in the lumen (Fig. 6) and that finally reach the reservoir where they are stored until the emergence (see the size of the reservoir in Fig. S22E).

The patterns of expression of all 13 genes fit our prediction: they are all specifically expressed in the venom glands of females but not in other tissues, nor in males (Fig. 4). Some virally-derived genes were particularly expressed at the very beginning of venom gland morphogenesis (day 11), whereas the other genes had their peak of expression at day 14, when the reservoir of the gland starts to be filled with VLPs.

Two sets of genes could also be identified base on their level of expression. One set of genes had an expression between 3 and 12 times that of the actin control gene (ORFs 94, 107, 60, 83 and 85), whereas the other genes had lower levels of expression, below 1.8 times that of the actin control (ORFs 5,72,68, 92, 87, 58, 78). ORF96 was even below the detection threshold in our assay.

Finally, we also measured the expression of a wasp virulence protein, known as a major component of wasp venom, most likely wrapped within the VLPs in *Leptopilina boulardi* (the RhoGAP LbGAP [43], [19], [28]). Contrary to the 13 virally-derived genes, this virulence protein has a eukaryotic origin[19]. As expected, this gene is also specifically expressed in the venom gland, and transcription starts just after the 14-day peak observed for most virally-derived genes. Interestingly, among “early” virally-derived genes, we identified a putative DNA polymerase (ORF58, see table 2). This opened the fascinating possibility that the DNA encoding those genes is amplified during this biological process.

**Table 2:**
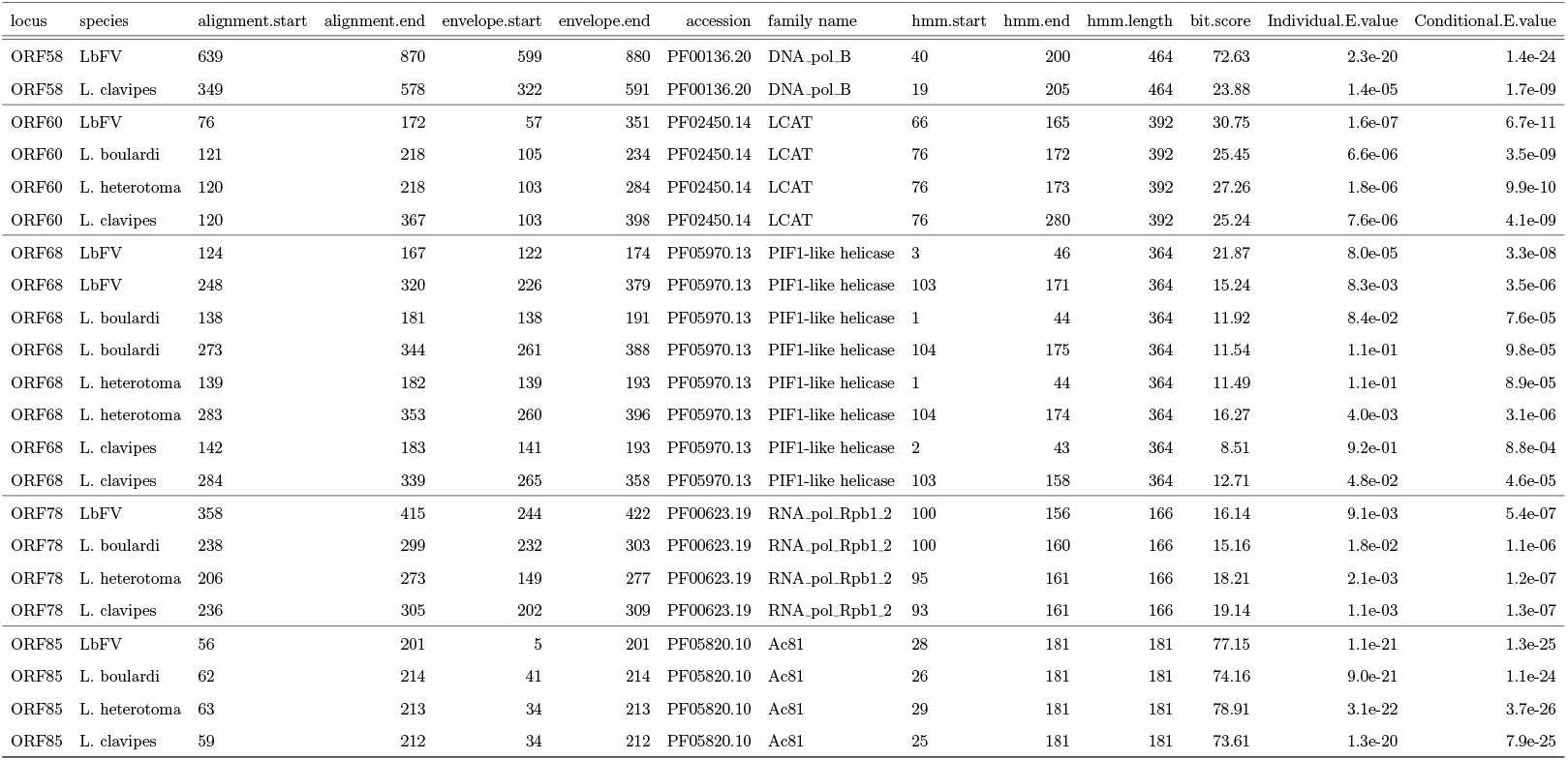
hmmer sequence analysis for the 13 proteins encoded by LbFV and their orthologs in *Leptopilina* wasps. Only hits with individual evalues < 0.15 are shown.

### 2.4 Most virally-derived genes but not the major wasp virulence factor are amplified in the venom gland

Using real-time PCR, we measured the relative DNA levels of each gene compared to an actin single copy locus. As in the transcription assay, we measured it in the venom gland, ovaries, rest of the body and in males of *L. boulardi*. We also included another single copy gene (shake) as a control. As expected the relative copy number of shake did not show any trend in time, nor differences between tissues, thus validating our assay (Fig. 5). We observed similar “flat” patterns for ORF87, ORF58 and ORF96 although a statistically significant effect was detected at day 11 for ORFs 87 and 96. On the contrary, all other virally-derived genes were significantly amplified in the venom gland, but not in other tissues. This amplification was highly significant for most genes at day 14, were they all reached their peak of amplification. Interestingly, among the three genes that were not amplified is the putative DNA-polymerase (ORF58). This gene showed an early-transcription profile in the transcriptomic assay. The same “early-gene expression pattern” is also observed for the other non-amplified gene (ORF87). For most virally-derived genes, we observed a striking correlation between the transcription and amplification profiles (compare figs. 4 and 5). Finally, our dataset indicates that the gene encoding the major constituent of VLPs (LbGAP) is not amplified (Fig. 5).

### 2.5 A virally-derived protein is present in mature VLPs of *Leptopilina sp*

In order to further test the hypothesis that the virally-derived genes are involved in VLP formation, we purified VLPs from adult *L. boulardi* females. Mass spectrometry-based proteomics was then used to identify proteins present in two independent biological replicates (fig S23). This strategy allowed the identification of a total of 383 proteins, of which 236 were found in both replicates. Among these proteins, as expected, we were able to reproducibly identify typical virulence proteins known to be part of VLP content (such as the RhoGap LbGAP [19], superoxide dismutase [17], serpin [16] or calreticulin [82]) confirming that we correctly purified the proteins (supplementary table S7). More importantly, in both biological samples we found the presence of the endogenized version of LbFV ORF85 protein (3 peptides in sample 1 and 2 in sample 2, supplementary table S7). Finally, we reanalyzed a similar proteomic dataset obtained by others [35] using the related species *L. heterotoma*. Again, we detected the endogenized version of LbFV ORF85 protein (although with a single peptide, data not shown). Taken together, these data demonstrate that the virally-derived protein ORF85 encoded in the genome of *Leptopilina* species is part of mature VLPs.

### 2.6 Annotation of virally-derived genes

Out of the 13 viral genes, five had similarities with known protein domains (table 2). First, the viral protein ORF58 showed clear similarity with DNA polymerase B domain (e-value 2.3 x 10^−20^). The domain was also detected in wasp orthologs but only for the *L. clavipes* protein. For the other four proteins, similar domains were identified in both the LbFV sequence and the wasp sequences. ORF60 bears a lecithine cholesterol acyl transferase (LCAT) domain, ORF68 contains a PIF1-like helicase, ORF78 contains an RNA-polymerase domain. Finally, ORF85, which is detected in mature VLPs, contains an Ac81 domain, a conserved protein found in all Baculoviruses [61], and known to be involved in virus envelopment [23].

## 3 Discussion

In this paper, we showed that all *Leptopilina* species contain a set of genes of viral origin deriving from either a direct ancestor of LbFV or from a closely related one. We describe the genomic structure of those genes in details in *L. boulardi, L. heterotoma* and *L. clavipes*, for which the whole genome was obtained. In addition, we were able to detect the presence of one LbFV-derived gene (ORF96) in all *Leptopilina* DNA extracts tested so far, suggesting that those virally-derived genes are shared by all *Leptopilina* species. Finally, one virally-derived protein (ORF85) is detected in purified VLPs. From this analysis, we conclude that an ancestor of all *Leptopilina* species acquired a set of 13 viral genes deriving from a virus related to the behavior manipulating virus LbFV. These genes have been conserved in all *Leptopilina* species and allow them to produce immuno-suppressive VLPs. This is very likely the consequence of a single event.

So far, all studied *Leptopilina* species are known to produce VLPs in their venom gland [68][57][32]. We confirmed this result in *L. boulardi* and found typical VLPs also in *L. clavipes*, suggesting that all *Leptopilina* species do produce VLPs. These particles are produced at the pupal stage and are stored in the reservoir of the venom gland. During oviposition, females inject not only their egg(s) but also some VLPs into their *Drosophila* hosts. VLPs are conceptually similar to liposomes that would contain virulence proteins. VLPs then permit the wasp to address these proteins to *Drosophila* immune cells [19]. The virulence proteins delivered to the target cells then induce important morphological changes in the lamellocytes, precluding them from initiating an efficient immune reaction against the parasitoid egg [19]. Thus, the VLPs are essential for the reproduction of the wasps. Because the proteins wrapped within the VLPs have a eukaryotic origin and because neither viral transcripts, viral proteins, nor viral DNA had been identified from venom gland analysis, it has been claimed that VLPs do not have a viral origin [66, 35]. In addition, the description of VLP proteins with eukaryotic microvesicular signature has been put forward as an evidence of a eukaryotic origin for these structures [35]. Following this argumentation, the authors proposed to change the denomination of VLPs for MSEV (mixed-strategy extracellular vesicle). On the contrary, our data strongly suggest that the VLPs found in *Leptopilina* do have a viral origin and derive from a massive endogenization event involving a virus related to an ancestor of the behaviour manipulating virus LbFV (Fig S2B). Under this scenario, present-day VLPs are indeed eukaryotic structures but evolved thanks to the endogenization and domestication of ancient viral genes. Nowadays, these structures allow the delivery of eukaryotic virulence proteins to *Drosophila* immune cells.

As expected from this hypothesis, we found that the virally-derived genes are specifically expressed in the venom gland, during the first part of the pupal stage, time at which the VLPs are beginning to be produced. In addition, those genes are under strong purifying selection, as could be expected for genes involved in the production of such fitness-related structures as VLPs. Analyzing the putative biological function of the genes brings additional support in favor of this hypothesis. Although 8 out of the 13 genes have no conserved domains, two of them have functions suggesting that they could be involved in membrane formation.

The first one is ORF60 which contains a lecithine cholesterol acyl transferase (LCAT) domain. In humans, LCAT is involved in extracellular metabolism of plasma lipoproteins, including cholesterol. LCAT esterifies the majority of free cholesterol, catalyzing translocation of fatty acid moiety of lecithin (phosphatidyl choline) to the free 3-OH group of cholesterol. It thus plays a major role in the maturation of HDL (high-density lipoprotein cholesterol) [69]. This putative biological property makes sense under our hypothesis since VLPs resemble liposomes that may be composed of highly hydrophobic compounds such as cholesterol. We may thus speculate that ORF60 plays a crucial role in the early formation of the “empty” membranes observed in the lumen of the venom gland under transmission electron microscopy (Fig. 2. 3A-B). Interestingly, the phylogenetic reconstruction of this gene suggests that LbFV itself acquired LCAT gene from a bacterial donor species.

**Figure 3:**
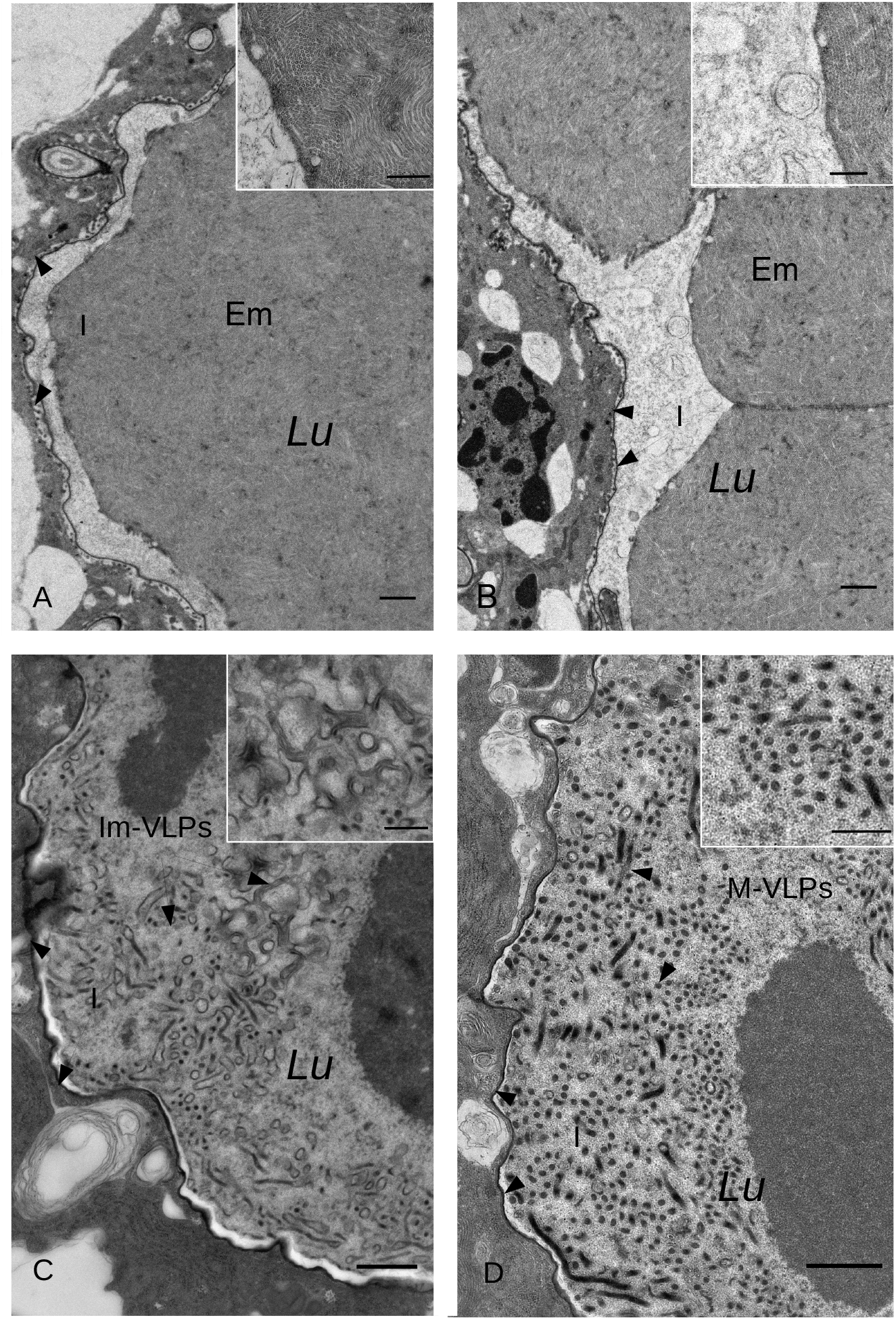
Biogenesis of VLPs in the venom gland of *L. boulardi* during the pupal stage until adult emergence: (A) 14 days (pupae), (B) 16 days (pupae), (C) 18 days (pupae), (D) 21 days (adult). At days 14 and 16, secretory cells (SC) are releasing empty membranes (Em) into the Lumen (Lu) of the venom gland where they accumulate. Then at day 18, empty membranes starts to be filled with electron-dense material (probably virulence proteins, such as LbGAP) to produce immature VLPs (im-VLPs). Finally at emergence (day 21), the venom gland lumen is filled with mature VLPs (m-VLPs) ready to be injected into the host. I: cuticular intima delineating the lumen. Inserts show details of each image. Bars represent 1*μ*M, except in inserts where they represent 500*μ*M.

The second relevant gene is ORF85. ORF85 is an homolog of Ac81, a conserved protein found in all Baculoviruses [61]. Its role has been recently deciphered in *Autographa californica* multiple nucleopolyhedrovirus (AcMNPV, [23]). During their cycle, baculoviruses first produce budded virions (BVs) and, late in infection, occlusion-derived virions (ODVs). After the initial infection, BVs are responsible for the spread of the infection from cell to cell within the infected insect. On the contrary, ODVs are only produced at the final stage of the infection. At that point nucleocapsids are retained in the nucleus where they acquire an envelope from microvesicles. They are then exported into the cytoplasm and are embedded into proteinaceous crystal matrix, thus forming occlusion bodies (OBs). The OBs are then released in the environment. OBs are absolutely necessary to initiate new insect infection through horizontal transmission. By a mutant analysis, Dong *et al*. [23] showed that Ac81 is necessary for the capsid envelopment and embedding within the occlusion bodies (OBs). They also showed that Ac81 contains an hydrophobic transmembrane domain that is necessary for this step. Interestingly, all three orthologs in *Leptopilina* sp. also contain a TM domain (Fig. S24). Our hypothesis is that the homolog of Ac81 in *Leptopilina* species is involved in the wrapping of virulence proteins into the VLPs, which is observed at day 18 under electron microscopy (Fig. 2.3C). Interestingly, it has been found that the closest viral homolog of this protein (apart from LbFV) is a structural protein of the Hytrosaviridae GpSGHV. In line with this, we found that protein ORF85 is indeed part of mature VLPs in *L. boulardi* and *L. heterotoma* and very likely in all *Leptopilina* species. This protein thus probably plays a crucial role in wrapping virulence proteins into VLP membranes and/or in the fusion with the target *Drosophila* immune cells. Interestingly, a nudiviral homolog of Ac81 has also been domesticated by *Venturia canescens* where three paralogs are found [47].

The other genes containing a conserved domain reveal functions related to DNA replication and transcription. The presence of a putative DNA polymerase (ORF58) and an helicase (ORF68) may sound surprising if one considers that VLPs do not contain DNA, contrary to polydnaviruses. However, we observed that after the early transcription activation of the DNA polymerase (at day 11), 10 out of the 13 virall-derived genes were subsequently amplified (at day 14). This genomic amplification correlates very well with their respective expression profile which suggests that the transcriptomic regulation of these virally-derived genes is governed, at least partly, by the gene copy number in the cell. Interestingly, the DNA polymerase itself and the nearby virally-derived gene (ORF87) are not amplified, suggesting that the amplification depends on the location of the loci in wasp chromosome. It is unclear at that point whether the genomic amplification involves the production of circular or linear amplicons or concatemers, and where are located the boundaries of the amplified loci. On the contrary, the gene encoding the major constituent of the VLPs (LbGAP), which does not have a viral-origin, is not genomically amplified, although it is highly transcribed from day 14 until the emergence of the wasp and finally detected in mature VLPs as a protein. This suggests that the virally-derived DNA polymerase targets some specific sequences flanking the amplified loci. The wasp genome also encodes a virally-derived RNA polymerase (ORF78) that is likely involved in the transcription of the virally-derived genes.

All together, our data strongly suggest that VLP production is possible thanks to the domestication of 13 virally-derived genes, captured from an ancestor of LbFV. Based on the clustering of the genes in *L. boulardi* assembly, and on the synteny conservation, we speculate that a single event led to the acquisition of the whole gene set. We can even hypothesize that a whole virus genome integrated into the chromosome of the *Leptopilina* ancestor. Several recent publications suggest that large, possibly full-genome insertions of symbiont into their host DNA do occur in the course of evolution, including from dsDNA viruses. For instance, whole genome sequencing of the brown planthopper revealed a total of 66 putative ORFs (74,730bp in total) deriving from a nudivirus genome, including 32 out of the 33 core nudiviral genes [13]. Also, it has been recently shown that an almost complete *Wolbachia* genome has been integrated into the chromosome of its host the common pillbug *Armadillidium vulgare*, with dramatic consequences on its sex determinism system[46]. After this suspected full-genome insertion of an ancestor of LbFV, we speculate that subsequent rearrangements have eliminated unnecessary genes and finally scattered, to a certain degree, the 13 remaining genes. Better genome assemblies are now necessary to gain insights on this aspect of the domestication process in the different *Leptopilina* lineages.

Our results document a novel domestication event of viruses in parasitic wasps. Indeed, from a function point of view, the domestication we document here is very similar to what has been described in the microgastroid complex in Braconidae[4], in Campopleginae and Banchinae [78][3] and in Opiinae [10]. In all cases, it is thought that a single endogenization event led to the integration of viral DNA into wasp chromosomes, and subsequently to the evolution of a virally-derived system delivering virulence factors to host immune cells. Despite these similarities, the underlying mechanisms are different. In the braconidae *Cotesia congregata* and *Microplitis demolitor* and in the Campopleginae *Hyposoter dydimator*, the putative virally-derived genes are genomically amplified as well as the genes encoding the virulence factors[50][11][78], although different mechanisms are involved[11]. The main consequence of this amplification is the production of the DNA circles that are finally packed into the polyDNAviruses.

On the contrary in *Leptopilina boulardi*, we find that only the 13 virally-derived genes are amplified, but not the virulence gene RhoGAP. The *Leptopilina* system best resembles the VLP production observed in *Venturia canescens* in the sense that VLP do not contain DNA (contrary to PolyDNAviruses described above) but instead proteins[35]. In *Leptopilina*, the genomic amplification seems to be an original trancriptional mechanism occurring during the production of the VLPs membranes. Virally-derived genes are also amplified during VLP production in *V. canescens* [47].

From these examples, it is clear that the domestication of whole sets of viral genes have repeatedly occurred in endoparasitoid wasps belonging to the super-family Ichneumonoidea, with at least two events leading to polydnavirus systems (that adress DNA circles encoding virulence factors to the host) in some Braconidae and Ichneumonidae and two events leading to the evolution of a VLP system (that address virulence proteins wrapped into a liposome-like structure to the host) in *Fopius species* (Opiinae) [10] and in *V. canescens* (Campopleginae) [36], [65]. Actually, this last VLP domestication in *V. canescens* better corresponds to a replacement of a PDV system by a VLP system[65], showing that domestication events have been frequent in this superfamily. With our results obtained on species belonging to the Figitidae family, which diverged from Ichneumonoidea 225My ago [64], it is tempting to extend this conclusion to other clades of Hymenoptera endoparasitoids. If this idea is confirmed, then a striking parallel comes up between virus domestication in Hymenoptera and syncytin domestication in mammals[45]. In both cases, viral proteins have been repeatedly co-opted to permit cell-cell fusion, although in one case this is for materno-fetal communication and in the second case it is for virulence factor delivery. Future investigations should test more thoroughly this hypothesis.

One remaining open question for all those events, is the type of interaction the ancestral virus and its wasp did have before the domestication happened. Regarding this question, very few data are available up to now. For PDV found in campopleginae such as *H. dydimator* and in banchinae such as *Glypta fumiferanae*), the ancestral virus has not been clearly identified[78][3]. On the contrary, the putative virus donors have been identified as a beta-nudivirus for PDVs in braconidae[4], and as an alpha-nudivirus for VLPs found in *Venturia canescens*[65] and in *Fopius* species[10]. However, their closest viral relatives are not infecting hymenoptera, but rather other arthropods[73]. In addition, the endogenization event is ancient, at least for Bracoviruses, which is the only case for which an estimation exists (103My, [60]), rendering difficult the inferences on the type of association that existed upon emergence of the association. It is thus unclear what type of interaction did the ancestral virus have with its host before the endogenization process.

In *Leptopilina*, we unequivocally identified an ancestor (or a close relative) of the behaviour-manipulating virus LbFV as the donor virus. First, it should be noted that in previous cases for which the ancestor has been identified the donor virus has a large circular genome composed of a double stranded DNA. Our results again show the same pattern. Second, the previous studies repeatedly identified nudiviruses as the donor family. Here we identify a virus belonging to another, possibly new, virus family[48]. This virus is related to nudiviruses and baculoviruses, but is more closely related to the hytrosaviruses [2], which are known to induce Salivary Gland Hypertrophy in tsetse flies and house flies, although it can also remain symptom-less [1].

Finally, this is the first time that the identified virus ancestor still has extant relatives infecting one of the wasp species. From our previous work on the interaction between LbFV and its host *Leptopilina boulardi*, we know that LbFV is vertically transmitted and replicate in cells of the oviduct[77]. This result suggests that physical proximity with the germ line may have facilitated the initial endogenization event, thus allowing the initiation of the domestication process. The identification of a contemporary virus still infecting the wasp also opens the way for addressing experimentally the mechanisms by which the virus could integrate into wasp chromosomes. Finally, LbFV is responsible for a behavior manipulation in *L. boulardi*: it forces females to superparasitize, which allows its horizontal transmission to other wasps[75]. This raises the fascinating possibility that the ancestral donor virus also manipulated the behavior of the wasp. To clarify this issue, the sampling of relatives of LbFV will be essential, to be able to reconstruct the ancestral state for the lineage that actually gave rise to such genetic innovation in wasp genomes.

## 4 Methods

### 4.1 Wasp rearing

*L. boulardi, L. heterotoma* and *G. brasiliensis* were reared on *D. melanogaster* as host (StFoy strain) in a climatic chamber (25C 60% humidity, 12/12 LD). The *G. brasiliensis* strain was kindly provided by Dr. Shubha Govind, *L. clavipes* by Dr. Elzemiek Geuverink and *L. boulardi* and *L. heterotoma* strains were collected and identified by our group. *Drosophila* were fed with a standard medium [22]. All experiments on *L. boulardi* were performed on a strain uninfected with the behaviour-manipulating virus (NSref).

### 4.2 Wasp genome sequences and annotation

We previously reported the genome of *Leptopilina boulardi*, strain Sienna (accession number : PQAT00000000) which has been obtained from the sequencing of a single female[76]. Although this female was infected by LbFV, the draft genome does not contain contigs belonging to the virus genome since we removed them by comparison to the published virus genome sequence[48]. The assembly was performed using IDBA_ud [63] followed by a scaffolding step with assembled RNAseq reads using the software L_RNA_scaffolder [81].

We sequenced the genomes of the related *L. heterotoma* (Gotheron strain, accession number RICB00000000), and the more distantly related *G. brasiliensis* (Va strain, accession number RJVV00000000). *L. heterotoma* is refractory to infection by LbFV[62] and no reads mapping to LbFV genome has been found neither in *L. heterotoma* nor in *G. brasiliensis* datasets. We extracted the DNA of a single female abdomen using Macherey-Nagel columns, similarly to what was performed for *L. boulardi* [76]. The DNAs were then used to prepare paired-end Illumina libraries using standard protocols (TruSeq PE Cluster v3, TruSeq SBS 200 cycles v3, TruSeq Multiplex Primer). The libraries were then sequenced on a Hiseq2500 (for L.h, 2 × 100bp, insert size = 418bp) or Hiseq3000 (for G.b, 2 × 150bp, insert size = 438bp) machine on the Genotoul sequencing platform.

Similarly to what was done for *L. boulardi*, the drafts of *L.heterotoma* and *G.brasiliensis* were obtained after assembling genomic DNA reads with IDBA_ud [63]. For *L. heterotoma* assembly, this was followed by scaffolding using publicly available assembled RNAseq reads[28] by running the software L_RNA_scaffolder[81]. This RNA-seq scaffolding step was not performed for *G. brasiliensis* because no RNAseq reads were available for this species in public databases.

The genome of an asexual strain of *L. clavipes* (strain GBW) which is not infected by LbFV was obtained and is described in [41] (accession PRJNA84205). To have comparable assembly strategies, we included an additional RNA scaffolding step using publicly available sequences ([56]).

In order to test the completeness of the drafts generated, we ran the BUSCO pipeline (version 2.0) that looks for the presence of 1066 ubiquitous genes shared by at least 90% of all arthropods ([71]).

The genome sizes were estimated using several methods. First of all, we simply divided the total number of bases mapped to the draft by the mean coverage observed on scaffolds containing complete BUSCO genes. Those scaffolds are expected to contain non repeated nuclear DNA and their coverage is a valuable estimate of the coverage for any nuclear locus. Second, after filtering out adapters containing reads with Skewer version 0.2.2[39], removing reads duplicates with FastUniq version 1. 1 [80], filtering out reads mapping to mitochondrial contigs with Bowtie 2 version 2.3.4.1[44] and samtools version 1.8[49], removing contaminant reads (from viruses, prokaryotes and microbial eukaryotes) with Kaiju 1.6.2 used with the NR+euk 2018-02-23 database[55], k-mers frequencies were established from the remaining reads for each species using Jellyfish 2.2.9[52] and k = 21 (default value). From these 21-mers distributions genome size was estimated with findGSE[72] used with default parameters. These estimates were then used to run DNAPipeTE version 1.3[30] (2 samples per run, 0.1X coverage per sample) in order to assess the repetitive fraction of the genomes. Finally, independant estimates from flow cytometry experiments were obtained for *L. boulardi, L. heterotoma* and *G. brasiliensis* from [29] and for *L. clavipes* from [41].

We predicted genes in wasp sequences using the software augustus 3.2.3 [37], with training parameters obtained from the BUSCO outputs.

### 4.3 Homology search

In order to identify homologies between viral proteins and wasp DNA, we used a simple tblastn (v. 2.6.0) approach with viral proteins as query and each wasp genome as database. Default parameters were used except that an evalue threshold of 0.01 was chosen.

### 4.4 Phylogenies

#### 4.4.1 Species-tree

Based on 627 “universal arthropod” genes identified by the BUSCO pipeline [71], a species tree was constructed for *L. heterotoma, L. boulardi, L. clavipes* and *G. brasiliensis*, using *Apis mellifera* as outgroup. The protein sequences were aligned using the bioconductor msa package[7]. Individual alignments were concatenated and a phylogenetic reconstruction was then performed using PhyML (parameters: -d aa -m LG -b -4 -v e -c 4 -a e -f m)[33]. In total, 290428 variable sites were found and the branch supports were computed using approximate likelihood ratio test (aLRT). We also constructed a tree for 10 *Leptopilina* species and *G. brasiliensis* using publicly available sequences of Internal transcribed spacer 2 (ITS2). Alignment was performed with muscle and a phylogeny was obtained with PhyML (parameters: -d nt -m GTR -b -4 -v 0.0 -c 4 -a e -f e). In total, 399 variable sites were used and the tree was rooted using mid-point rooting method.

#### 4.4.2 Gene-tree

We searched orthologs of viral proteins of interest in other organisms by blasting (blastp) them against nr (downloaded on october 2017) with an evalue threshold of 0.01. After retrieving the sequences, we selected one sequence per species and added them to the proteins identified in *Leptopilina* genomes. The sequences were then aligned using muscle algorithm v3.8.31. Because the proteins included in the alignment diverged considerably, we selected blocks of conserved sites using the gblocks algorithm parametrized with less stringent options (allowing smaller final blocks, gaps within final blocks and less strict flanking positions, [12]). Phylogenetic reconstruction was then performed using PhyML (parameters: -d aa -m LG -b -4 -v e -c 4 -a e -f m). The branch supports were computed using approximate likelihood ratio test (aLRT). The accession numbers of the sequences used in the phylogenies are reported in table S5.

### 4.5 PCR amplification of ORF96

Based on the sequences of *L. boulardi, L. heterotoma* and *L. clavipes*, we designed primers for the orthologs of LbFVORF96. The primer sequences are ATTGGTGAAATTCAATCGTC and TCATTCATTCGCAATAATTGTG. They amplified a 411bp internal fragment of the coding sequence. PCR reaction was performed in a 25uL volume containing 0.2uM primers, 0.2mM dNTPs, 1mM MgCl2 and 0.5U of Taq DNA polymerase with the following cycling conditions : 95 °C 30”, 54 °C 30”, 72 °C 60” (33 cycles).

### 4.6 dN/dS calculation

The coding sequences of “universal arthropod” BUSCO genes identified in the three *Leptopilina* species were extracted and, using the msa and seqinr R package, were reverse-aligned using the protein alignments as a guide (reverse.align function of the seqinr package). dN/dS ratios were then estimated using the kaks function of the seqinr R package. The method implemented in this package is noted LWL85 in [74]. A similar procedure was performed for the 13 virally-derived genes found in the genomes of the three *Leptopilina* species.

### 4.7 Expression in the venom gland and other tissues

We studied the expression of genes during the pupal stage of *L. boulardi*, at days 11, 14, 16, 18 and 21. The wasp strain used is not infected by the behaviour-manipulating virus LbFV. 11 days corresponds to the beginining of the pupal stage, whereas 21 days corresponds to the emergence time. Wasps were gently extirpated from the *Drosophila* puparium, and venom gland, ovaries, rest of the body of *L. boulardi* females was dissected in a droplet of PBS + 0.01% tween and deposited in the RLT+B-mercaptoethanol buffer of the Qiagen RNAeasy extraction kit. Males were also prepared as a control, in a similar way. The tissues extracted from twenty individuals were then pooled together and tissues were disrupted in a Qiagen homogenizer (3 minutes 25Hz). Two biological replicates were performed for each condition, except for day 11 where only one sample was obtained. cDNAs were synthetized using the SuperscriptIII kit (ThermoFisher). Real-time PCR assays were then performed with SYBR green (ssoadvanced universal sybr green supermix, Biorad) using standard procedures on a Biorad CFX-96 machine. We quantified the number of copies of each target cDNA using a serial dilution standards. Because we obtained only tiny quantities of RNA from this experiment (because of the very small size of the tissues dissected), we were not able to test numerous genes. We thus choose to use only one control gene (actin gene). As a counterpart, we were able to test all thirteen virally-derived genes and the RhoGAP gene. The primer sequences are given in table S6.

### 4.8 Genomic Amplification

Using a similar assay, we extracted the DNA of *L. boulardi*, at days 11, 14, 16, 18 and 21, using an uninfected strain (no LbFV present). The genomic DNA of 15 pooled individuals was extracted using the Nucleospin tissue Macherey-Nagel kit following provider’s instructions. Three biological replicates per condition was done. Real-time PCR assays were then performed with SYBR green using standard procedures on a Biorad CFX-96 machine. We quantified the number of copies of each target genes using a serial dilution standards. The primer sequences are given in table S1. For an unknown reason, the amplification with DNA extracted from ovaries was particularly difficult, in particular when the ovaries were mature (at day 21). We thus had to remove this tissue from the statistical analysis because Cqs were too high to be reliable. For the same reason, most data for ovaries at day 21 were removed from figure 5. The primer sequences are given in table S6. Shake and actin genes were chosen as single copy genes. This was checked by looking at the blast results using each primer set (a single 100% match was observed for both pairs of primers). Accordingly, a single band of the expected size was observed on a gel and the expected sequence was obtained after Sanger-sequencing for both loci.

### 4.9 Statistical analysis

For both the transcriptomic and genomic analysis, we calculated the absolute copy number of each gene of interest and divided it by the absolute copy number of the actin control gene. This ratio was then analyzed in an anova framework with time, tissue and time:tissue interation as factors. The effects were tested by likelihood ratio tests (LRT) of full model versus reduced one. Contrasts between tissues were also calculated at each time point (corresponding to the star in figures 4 and 5). Residuals of the models were judged as unstructured and had an overall normal distribution.

**Figure 4:**
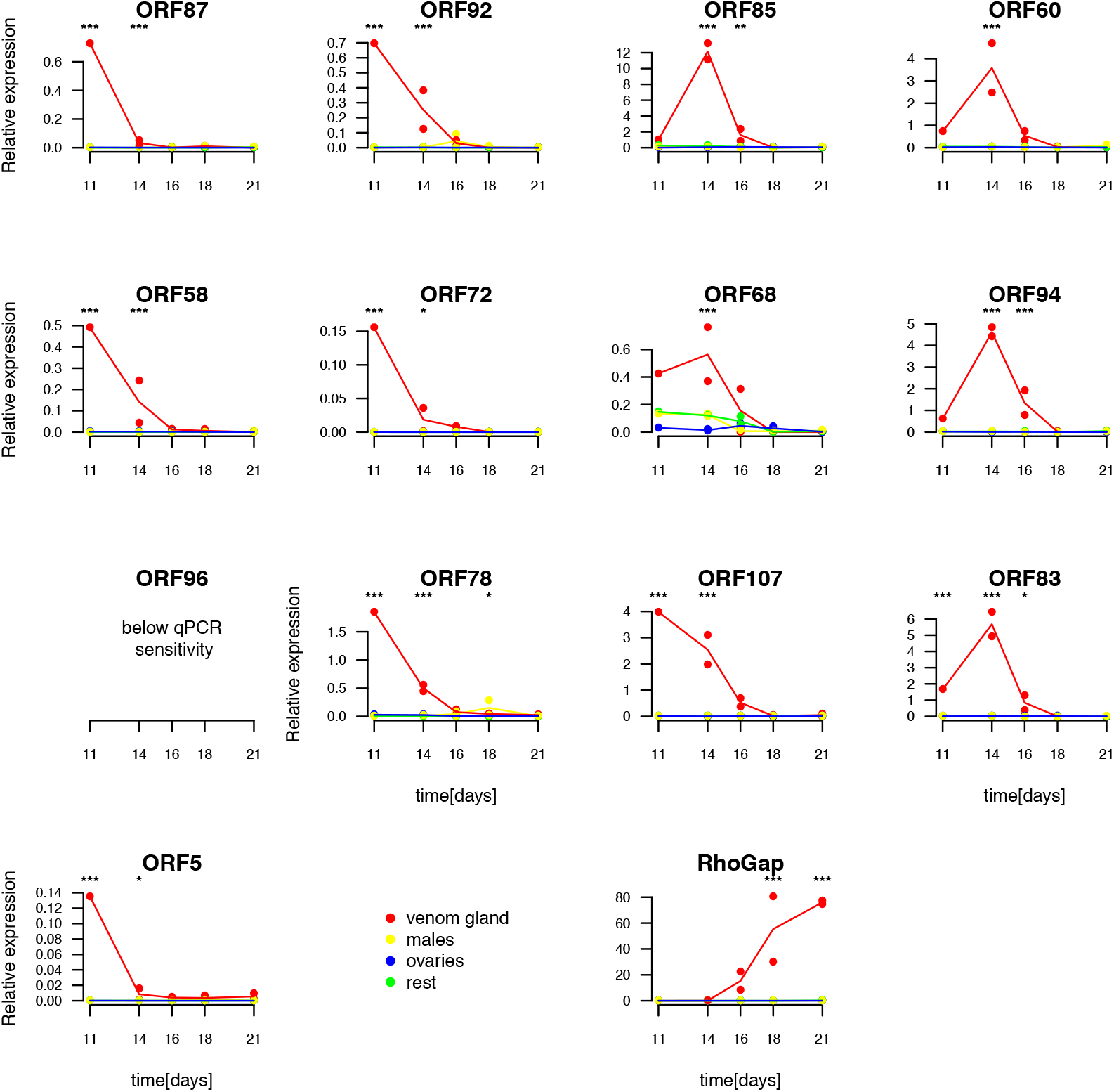
Expression of the 13 virally-derived genes and of the Rho-Gap in different tissues of *L. boulardi* from initial pupal stage to adult. x-axis represents days since egg-laying. 11 days corresponds to the beginning of the pupal stage and 21 days to the emergence of adults from the *Drosophila* puparium.

**Figure 5:**
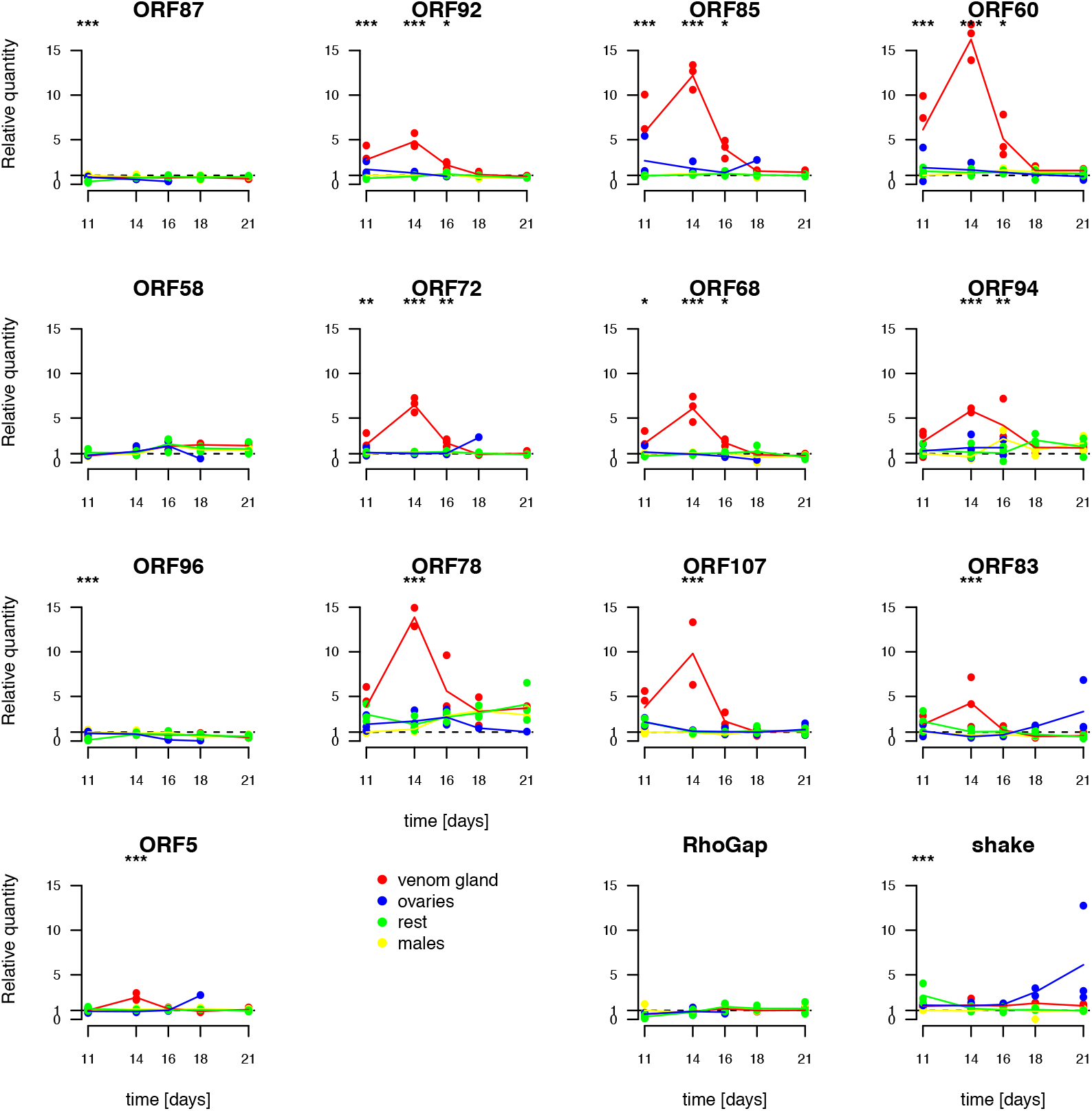
Genomic amplification of virally derived-genes measured by real time PCR in *L. boulardi*. The relative quantity of each target gene is represented relative to the actin control gene and normalized by the ratio observed in males at day 11. The expected value under no amplification (relative quantity=1) is indicated as a dotted line. Stars correspond to the tissue effect tested at each time point (with holm correction for multiple tests) : * < 0.05, ** < 0.01, *** < 0.001.

### 4.10 Morphogenesis and electron microscopy of the venom gland

To follow the morphogenesis of the venom gland, we dissected *L. boulardi* pupae at days 11, 14, 16, 18 and 21, in a similar design used for transcriptomics. Wasps were gently extirpated from the *Drosophila* puparium, and the venom gland of females was dissected in a droplet of PBS + 0.01% tween. Venom glands were either directly mounted on a glass slide for further examination under a light microscope or transfered into a solution of 2% glutaraldehyde in PBS for further examination under the Transmission Electron Microscope (TEM). For TEM, the tissues were then post fixed 1 hour in 2% osmium tetroxide in the same buffer, thoroughly rinced in distilled water, stained “en bloc” with a 5% aqueous uranyl acetate solution, dehydrated in a series of graded ethanol and embedded in Epon’s medium. Ultrathin sections were cut on a LKB ultratome and double stained in Uranyless and lead citrate. Samples were examined with a Jeol 1200 Ex transmission microscope at 80kV. Images were taken with an Quemesa 11 megapixel Olympus camera and analyzed with ImageJ software (https://imagej.nih.gov/ij/).

### 4.11 Proteomics

Proteins extracted from purified VLPs were using Laemmli buffer were stacked in the top of a SDS-PAGE gel (4-12% NuPAGE, Life Technologies), stained with Coomassie blue R-250 and in-gel digested using modified trypsin (Promega, sequencing grade) as previously described[70]. Resulting peptides were analyzed by online nanoliquid chromatography coupled to tandem mass spectrometry (UltiMate 3000 RSLC nano and Q-Exactive HF, Thermo Scientific). Peptides were sampled on a 300 *μ*m × 5 mm PepMap C18 precolumn and separated on a 75 *μ*m × 250 mm C18 column (Reprosil-Pur 120 C18-AQ, 1.9 *μ*m, Dr. Maisch) using a 120-min gradient. MS and MS/MS data were acquired using Xcalibur (Thermo Scientific). Peptides and proteins were identified using Mascot (version 2.6) through concomitant searches against the homemade *L. boulardi* database (see 4.2 for details), classical contaminant database and the corresponding reversed databases. The Proline software (http://proline.profiproteomics.fr) was used to filter the results: conservation of rank 1 peptides, peptide identification false discovery rate < 1% as calculated on peptide scores by employing the reverse database strategy and minimum of 1 specific peptide per identified protein group. Proline was then used to perform a compilation, grouping and spectral counting-based comparison of the protein groups identified in the different samples. Proteins from the contaminant database were discarded from the final list of identified proteins.

### 4.12 Annotation of viral genes

We searched for the presence of conserved domains in the 13 LbFV proteins horizontally transfered to *Leptopilina* species using the hmmer webserver (https://www.ebi.ac.uk/Tools/hmmer/) accessed the 5 of may 2018.

## 5 Acknowledgment

This work was supported by a grant from the Agence Nationale de la Recherche (ANR) to JV (11-JSV7-0011 Viromics). The bio-informatic work was performed using the computing facilities of the CC LBBE/PRABI. We thank the labex Ecofect for financial support for the internship of DDG. Proteomic experiments were partly supported by the Agence Nationale de la Recherche (ProFI grant ANR-10-INBSQ6). We thank Shubha Govind for kindly providing the *Ganaspis brasiliensis* strain, Elzemiek Geuverink for providing *L. clavipes* samples and Ken Kraaijeveld for providing access to raw Illumina reads of *L. clavipes*. We thank Dominique Colinet and Nathan Mortimer for sharing mass-spec data and analysis. We thank B. Bennetot for helpful preliminary data generation, and D. Kahn for advices on data analysis. We thank PCI Evol Biol reviewers for helpful comments on an earlier draft which has been reviewed and recommended by Peer Community In Evolutionary Biology (https://dx.doi.org/10.24072/pci.evolbiol.100062). Scripts used for this publication are available at https://doi.org/10.5281/zenodo.1889392.

