## supplementary tables and figures for "A behavior-manipulating virus relative as a source of adaptive genes for parasitoid wasps"

### 7 Supplementary tables and figures

| | basic statistics | | | | BUSCO stats | | | | | | Genome size [Mb] | | |
|---|---|---|---|---|---|---|---|---|---|---|---|---|---|
| species | n_scaffolds | N50 | coverage | Repetitive | Complete | Duplicated | Fragmented | Missing | total | missing | BUSCO.based | kmer.based | Cytometry.based |
| *L. boulardi* | 127707 | 14511 | 46 | 27.65% | 1044 | 4 | 8 | 10 | 1066 | 1% | 353 | 347 | 361 |
| *L. heterotoma* | 231242 | 12807 | 53 | 28.82 % | 1041 | 2 | 9 | 14 | 1066 | 1% | 445 | 464 | 459 |
| *L. clavipes* | 38495 | 17657 | 83 | 24.02 % | 1025 | 7 | 15 | 19 | 1066 | 2% | 257 | 300 | 321 |
| *G. brasiliensis* | 2777766 | 2080 | 24 | 44.92 % | 830 | 8 | 192 | 36 | 1066 | 3% | 829 | 977 | 968 |

Table S1: Statistics for the assemblies of wasp genomes. Genome size was estimated either using the coverage on BUSCO gene containing scaffolds or using a k-mer approach. For comparison, we give the estimated genome sizes obtained from flow cytometry analysis [29][41].

| | | | blast output | | | | | | | | | corresponding ORF on scaffold | | | | scaffold statistics | | |
|---|---|---|---|---|---|---|---|---|---|---|---|---|---|---|---|---|---|---|
| | query_id | query_len | subject_id | identity | aln length | qstart | qend | sstart | send | evalue | bitscore | start | end | length | strand | scaf_length | cov_depth | GC |
| 1 | LbFV_ORF5 | 696 | scaffold_159 | 34.4 | 366 | 337 | 696 | 6401 | 5337 | 5.5e-41 | 164.00 | 7601 | 5337 | 755 | + | 435056 | 53 | 0.36 |
| 2 | LbFV_ORF72 | 106 | scaffold_159 | 31.8 | 107 | 2 | 102 | 88433 | 88753 | 5.2e-10 | 57.40 | 88025 | 88771 | 249 | - | 435056 | 53 | 0.36 |
| 3 | LbFV_ORF92 | 1593 | scaffold_159 | 33.8 | 1058 | 583 | 1593 | 91842 | 94901 | 2.9e-151 | 518.00 | 89832 | 94901 | 1690 | - | 435056 | 53 | 0.36 |
| 4 | LbFV_ORF107 | 625 | scaffold_159 | 29.8 | 322 | 320 | 625 | 96312 | 95377 | 1.3e-11 | 71.20 | 97248 | 95377 | 624 | + | 435056 | 53 | 0.36 |
| 5 | LbFV_ORF94 | 182 | scaffold_159 | 29.0 | 176 | 1 | 173 | 98066 | 98557 | 5.5e-14 | 72.00 | 97829 | 98569 | 247 | - | 435056 | 53 | 0.36 |
| 6 | LbFV_ORF68 | 645 | scaffold_159 | 34.1 | 646 | 29 | 642 | 150985 | 152847 | 6.7e-99 | 335.00 | 150889 | 152856 | 656 | - | 435056 | 53 | 0.36 |
| 7 | LbFV_ORF60 | 362 | scaffold_159 | 32.6 | 377 | 5 | 353 | 187445 | 186375 | 2.4e-36 | 143.00 | 187532 | 186366 | 389 | + | 435056 | 53 | 0.36 |
| 8 | LbFV_ORF85 | 215 | scaffold_159 | 36.4 | 225 | 1 | 212 | 190829 | 190170 | 3.0e-26 | 108.00 | 190829 | 190149 | 227 | + | 435056 | 53 | 0.36 |
| 9 | LbFV_ORF87 | 176 | scaffold_2503 | 30.9 | 162 | 8 | 158 | 8659 | 8183 | 6.5e-12 | 65.90 | 8698 | 8078 | 207 | + | 55139 | 44 | 0.22 |
| 10 | LbFV_ORF58 | 1308 | scaffold_2503 | 36.7 | 932 | 3 | 904 | 10711 | 13299 | 1.3e-129 | 446.00 | 10909 | 14550 | 1214 | - | 55139 | 44 | 0.22 |
| 11 | LbFV_ORF78 | 676 | IDBA_scaffold_13958 | 40.1 | 670 | 43 | 670 | 2268 | 4205 | 1.2e-134 | 434.00 | 2487 | 4241 | 585 | - | 4800 | 49 | 0.57 |
| 12 | LbFV_ORF83 | 433 | scaffold_2315 | 24.8 | 435 | 14 | 407 | 874 | 2139 | 1.6e-15 | 82.40 | 862 | 2259 | 466 | - | 22591 | 45 | 0.20 |
| 13 | LbFV_ORF96 | 1048 | IDBA_scaffold_2184 | 41.9 | 1024 | 48 | 1041 | 3609 | 6512 | 4.0e-169 | 554.00 | 3564 | 6545 | 994 | - | 14197 | 45 | 0.28 |

Table S2: Blast hits for the 13 viral genes against *L. boulardi* genome.

| | | | blast output | | | | | | | | | corresponding ORF on scaffold | | | | scaffold statistics | | |
|---|---|---|---|---|---|---|---|---|---|---|---|---|---|---|---|---|---|---|
| | query_id | query_len | subject_id | identity | aln length | qstart | qend | sstart | send | evalue | bitscore | start | end | length | strand | scaf_length | cov_depth | GC |
| 1 | LbFV_ORF5 | 696 | IDBA_scaffold_8257 | 29.7 | 370 | 333 | 696 | 6582 | 7661 | 3e-37 | 157.00 | 5424 | 7661 | 746 | - | 9987 | 59 | 0.29 |
| 2 | LbFV_ORF72 | 106 | IDBA_scaffold_32827 | 28.6 | 70 | 34 | 102 | 1541 | 1750 | 4e-04 | 36.60 | 1303 | 1563 | 87 | - | 2607 | 58 | 0.23 |
| 3 | LbFV_ORF92 | 1593 | IDBA_scaffold_7081 | 38.1 | 501 | 1109 | 1590 | 5437 | 3938 | 5e-94 | 347.00 | 9070 | 3929 | 1714 | + | 10934 | 53 | 0.29 |
| 4 | LbFV_ORF107 | 625 | IDBA_scaffold_7081 | 27.1 | 170 | 455 | 621 | 2550 | 3056 | 9e-09 | 62.40 | 1179 | 3065 | 629 | - | 10934 | 53 | 0.29 |
| 5 | LbFV_ORF94 | 182 | IDBA_scaffold_13988 | 27.6 | 174 | 1 | 171 | 2671 | 2186 | 1e-11 | 69.70 | 2905 | 2168 | 246 | + | 5494 | 53 | 0.23 |
| 6 | LbFV_ORF68 | 645 | IDBA_scaffold_6001 | 32.6 | 660 | 29 | 644 | 7459 | 5555 | 3e-92 | 339.00 | 7561 | 5552 | 670 | + | 11133 | 52 | 0.48 |
| 7 | LbFV_ORF60 | 362 | scaffold_1324 | 26.0 | 381 | 5 | 353 | 4186 | 3065 | 7e-30 | 131.00 | 4270 | 3056 | 405 | + | 11549 | 50 | 0.34 |
| 8 | LbFV_ORF85 | 215 | scaffold_1324 | 35.2 | 219 | 1 | 207 | 375 | 1031 | 1e-23 | 109.00 | 375 | 1052 | 226 | - | 11549 | 50 | 0.34 |
| 9 | LbFV_ORF87 | 176 | IDBA_scaffold_5653 | 29.0 | 162 | 8 | 162 | 5879 | 6355 | 1e-05 | 49.70 | 5834 | 6457 | 208 | - | 11655 | 53 | 0.32 |
| 10 | LbFV_ORF58 | 1308 | IDBA_scaffold_5653 | 31.5 | 1378 | 19 | 1299 | 5204 | 1260 | 8e-158 | 558.00 | 5126 | 1170 | 1319 | + | 11655 | 53 | 0.32 |
| 11 | LbFV_ORF78 | 676 | IDBA_scaffold_9791 | 41.0 | 646 | 70 | 669 | 3914 | 2034 | 2e-123 | 443.00 | 3692 | 1992 | 567 | + | 9362 | 52 | 0.21 |
| 12 | LbFV_ORF83 | 433 | IDBA_scaffold_9791 | 21.9 | 429 | 14 | 407 | 7018 | 8277 | 8e-15 | 82.00 | 7006 | 8385 | 460 | - | 9362 | 52 | 0.21 |
| 13 | LbFV_ORF96 | 1048 | IDBA_scaffold_1712 | 36.6 | 1043 | 48 | 1041 | 16775 | 13806 | 2e-164 | 580.00 | 16820 | 13773 | 1016 | + | 26871 | 53 | 0.29 |

Table S3: Blast hits for the 13 viral genes against *L. heterotoma* genome.

| | | | blast output | | | | | | | | | corresponding ORF on scaffold | | | | scaffold statistics | | |
|---|---|---|---|---|---|---|---|---|---|---|---|---|---|---|---|---|---|---|
| | query_id | query_len | subject_id | identity | aln length | qstart | qend | sstart | send | evalue | bitscore | start | end | length | strand | scaf_length | cov_depth | GC |
| 1 | LbFV_ORF5 | 696 | scf7180005159507 | 33.1 | 366 | 337 | 696 | 1730 | 663 | 1.9e-40 | 162.00 | 2906 | 663 | 748 | + | 5318 | 87 | 0.31 |
| 2 | LbFV_ORF72 | 106 | scf7180005166731 | 32.7 | 107 | 2 | 102 | 6537 | 6217 | 8.8e-09 | 53.90 | 6945 | 6199 | 249 | + | 8832 | 81 | 0.30 |
| 3 | LbFV_ORF92 | 1593 | scaffold_1017 | 33.7 | 998 | 579 | 1536 | 21309 | 18403 | 3.1e-136 | 472.00 | 23376 | 18370 | 1669 | + | 23961 | 75 | 0.27 |
| 4 | LbFV_ORF107 | 625 | scf7180005156365 | 28.3 | 378 | 265 | 622 | 1897 | 809 | 5.3e-10 | 65.50 | 2674 | 803 | 624 | + | 5122 | 96 | 0.30 |
| 5 | LbFV_ORF94 | 182 | scf7180005161552 | 27.0 | 174 | 1 | 171 | 2763 | 2278 | 1.2e-12 | 67.80 | 3015 | 2260 | 252 | + | 4524 | 62 | 0.28 |
| 6 | LbFV_ORF68 | 645 | scf7180005174277 | 34.0 | 674 | 29 | 644 | 5118 | 7034 | 3.5e-103 | 347.00 | 5016 | 7037 | 674 | - | 7741 | 213 | 0.30 |
| 7 | LbFV_ORF60 | 362 | scf7180005174113 | 31.8 | 384 | 5 | 353 | 2297 | 3421 | 1.4e-33 | 134.00 | 2213 | 3430 | 406 | - | 6683 | 57 | 0.29 |
| 8 | LbFV_ORF85 | 215 | scf7180005171671 | 33.0 | 218 | 1 | 207 | 3017 | 3670 | 1.3e-23 | 100.00 | 3017 | 3691 | 225 | - | 4425 | 83 | 0.29 |
| 9 | LbFV_ORF87 | 176 | scaffold_886 | 31.5 | 165 | 8 | 158 | 8088 | 8570 | 3.6e-11 | 63.20 | 8049 | 8678 | 210 | - | 19330 | 85 | 0.28 |
| 10 | LbFV_ORF58 | 1308 | scf7180005154334 | 31.5 | 1042 | 317 | 1288 | 16626 | 13723 | 1.8e-120 | 418.00 | 16746 | 13633 | 1038 | + | 16768 | 70 | 0.26 |
| 11 | LbFV_ORF78 | 676 | scf7180005177077 | 41.0 | 675 | 39 | 669 | 11274 | 13268 | 3.7e-135 | 441.00 | 11517 | 13316 | 600 | - | 21465 | 86 | 0.29 |
| 12 | LbFV_ORF83 | 433 | scf7180005174071 | 24.5 | 436 | 9 | 404 | 3734 | 5005 | 1.8e-20 | 97.40 | 3740 | 5122 | 461 | - | 13231 | 85 | 0.28 |
| 13 | LbFV_ORF96 | 1048 | scf7180005173345 | 40.4 | 1013 | 48 | 1021 | 9667 | 6782 | 1.3e-178 | 582.00 | 9712 | 6686 | 1009 | + | 24926 | 74 | 0.28 |

Table S4: Blast hits for the 13 viral genes against *L. clavipes* genome.

Table S5: Accession numbers of sequences used in the phylogenies

| | Locus | species | GI | Figure |
|---|---|---|---|---|
| 1 | ORF5 | Lb | PQAT00000000 | 2 |
| 2 | ORF5 | Lh | RICB00000000 | 2 |
| 3 | ORF5 | Lc | JUFY01000000 | 2 |
| 4 | ORF5 | LbFV | 1148998810 | 2 |
| 5 | ORF58 | Lb | PQAT00000000 | 2 |
| 6 | ORF58 | Lh | RICB00000000 | 2 |
| 7 | ORF58 | Lc | JUFY01000000 | 2 |
| 8 | ORF58 | LbFV | 1148998708 | 2 |
| 9 | ORF60 | Lb | PQAT00000000 | 2 |
| 10 | ORF60 | Lh | RICB00000000 | 2 |
| 11 | ORF60 | Lc | JUFY01000000 | 2 |
| 12 | ORF60 | Lymphocystis_disease_virus_-_isolate_China | 51870153 | 2 |
| 13 | ORF60 | Organic_Lake_phycodnavirus_1 | 222510829 | 2 |
| 14 | ORF60 | Invertebrate_iridovirus_25 | 589287870 | 2 |
| 15 | ORF60 | Lymphocystis_disease_virus_1 | 611962711 | 2 |
| 16 | ORF60 | Lymphocystis_disease_virus_Sa | 1135106808 | 2 |
| 17 | ORF60 | LbFV | 1148998761 | 2 |
| 18 | ORF68 | Acyrthosiphon_pisum | 228698707 | 2 |
| 19 | ORF68 | Adoxophyes_honmai_entomopoxvirus_L | 506498063 | 2 |
| 20 | ORF68 | Apis_cerana_cerana | 1241837182 | 2 |
| 21 | ORF68 | Apis_dorsata | 572314547 | 2 |
| 22 | ORF68 | Apis_florea | 820863019 | 2 |
| 23 | ORF68 | Apis_mellifera | 571506210 | 2 |
| 24 | ORF68 | Bombus_terrestris | 240708910 | 2 |
| 25 | ORF68 | Camponotus_floridanus | 752871224 | 2 |
| 26 | ORF68 | Cephus_cinctus | 1000753753 | 2 |
| 27 | ORF68 | Chlamydotis_macqueenii | 677160893 | 2 |
| 28 | ORF68 | Crassostrea_gigas | 1139814932 | 2 |
| 29 | ORF68 | Cuculus_canorus | 676590237 | 2 |
| 30 | ORF68 | Dendroctonus_ponderosae | 546685733 | 2 |
| 31 | ORF68 | Diaphorina_citri | 662192917 | 2 |
| 32 | ORF68 | Diuraphis_noxia | 985403395 | 2 |
| 33 | ORF68 | Dufourea_novaeangliae | 987914045 | 2 |
| 34 | ORF68 | Eufriesea_mexicana | 1059214553 | 2 |
| 35 | ORF68 | Glossina_morsitans_morsitans | 83595237 | 2 |
| 36 | ORF68 | Gb | RJVV00000000 | 2 |
| 37 | ORF68 | Habropoda_laboriosa | 1059864473 | 2 |
| 38 | ORF68 | Harpegnathos_saltator | 749795708 | 2 |
| 39 | ORF68 | Helicoverpa_armigera | 204423112 | 2 |
| 40 | ORF68 | Lasius_niger | 861651735 | 2 |
| 41 | ORF68 | Lb | PQAT00000000 | 2 |
| 42 | ORF68 | LbFV | 1148998769 | 2 |
| 43 | ORF68 | Lc | JUFY01000000 | 2 |
| 44 | ORF68 | Lh | RICB00000000 | 2 |
| 45 | ORF68 | Myzus_persicae | 1230193237 | 2 |
| 46 | ORF68 | Nasonia_vitripennis | 1032757220 | 2 |
| 47 | ORF68 | Opisthocomus_hoazin | 677549512 | 2 |
| 48 | ORF68 | Papilio_machaon | 930680047 | 2 |
| 49 | ORF68 | Papilio_xuthus | 910339325 | 2 |
| 50 | ORF68 | Parasteatoda_tepidariorum | 1009572498 | 2 |
| 51 | ORF68 | Pogonomyrmex_barbatus | 769838565 | 2 |
| 52 | ORF68 | Polistes_canadensis | 954577453 | 2 |
| 53 | ORF68 | Trichomalopsis_sarcophagae | 1227108847 | 2 |
| 54 | ORF68 | Trichoplusia_ni | 6635437 | 2 |
| 55 | ORF68 | Vollenhovia_emeryi | 795079157 | 2 |
| 56 | ORF72 | Lb | PQAT00000000 | 2 |
| 57 | ORF72 | Lh | RICB00000000 | 2 |
| 58 | ORF72 | Lc | JUFY01000000 | 2 |
| 59 | ORF72 | Glossina_pallidipes_salivary_gland_hypertrophy_virus | 168804090 | 2 |
| 60 | ORF72 | LbFV | 1148998771 | 2 |
| 61 | ORF78 | Lb | PQAT00000000 | 2 |
| 62 | ORF78 | Lh | RICB00000000 | 2 |
| 63 | ORF78 | Lc | JUFY01000000 | 2 |
| 64 | ORF78 | LbFV | 1148998775 | 2 |
| 65 | ORF83 | Lb | PQAT00000000 | 2 |
| 66 | ORF83 | Lh | RICB00000000 | 2 |
| 67 | ORF83 | Lc | JUFY01000000 | 2 |
| 68 | ORF83 | Musca_domestica_salivary_gland_hypertrophy_virus | 187903111 | 2 |
| 69 | ORF83 | Glossina_pallidipes_salivary_gland_hypertrophy_virus | 984290647 | 2 |
| 70 | ORF83 | Glossina_pallidipes_salivary_gland_hypertrophy_virus | 984290648 | 2 |
| 71 | ORF83 | LbFV | 1148998781 | 2 |
| 72 | ORF85 | Lb | PQAT00000000 | 2 |
| 73 | ORF85 | Lh | RICB00000000 | 2 |
| 74 | ORF85 | Lc | JUFY01000000 | 2 |
| 75 | ORF85 | LbFV | 1148998786 | 2 |
| 76 | ORF87 | Lb | PQAT00000000 | 2 |
| 77 | ORF87 | Lh | RICB00000000 | 2 |
| 78 | ORF87 | Lc | JUFY01000000 | 2 |
| 79 | ORF87 | Phthorimaea_operculella_granulovirus | 21686761 | 2 |
| 80 | ORF87 | Agrotis_segetum_granulovirus | 46309360 | 2 |
| 81 | ORF87 | Spodoptera_litura_granulovirus | 148368915 | 2 |
| 82 | ORF87 | Glossina_pallidipes_salivary_gland_hypertrophy_virus | 168804094 | 2 |
| 83 | ORF87 | Musca_domestica_salivary_gland_hypertrophy_virus | 187903145 | 2 |
| 84 | ORF87 | Hemileuca_sp._nucleopolyhedrovirus | 529218126 | 2 |
| 85 | ORF87 | Spodoptera_frugiperda_granulovirus | 761719624 | 2 |
| 86 | ORF87 | Sucra_jujuba_nucleopolyhedrovirus | 960494866 | 2 |
| 87 | ORF87 | Glossina_pallidipes_salivary_gland_hypertrophy_virus | 984290700 | 2 |
| 88 | ORF87 | LbFV | 1148998788 | 2 |
| 89 | ORF92 | Lb | PQAT00000000 | 2 |
| 90 | ORF92 | Lh | RICB00000000 | 2 |
| 91 | ORF92 | Lc | JUFY01000000 | 2 |
| 92 | ORF92 | LbFV | 1148998790 | 2 |
| 93 | ORF94 | Lb | PQAT00000000 | 2 |
| 94 | ORF94 | Lh | RICB00000000 | 2 |
| 95 | ORF94 | Lc | JUFY01000000 | 2 |
| 96 | ORF94 | Glossina_pallidipes_salivary_gland_hypertrophy_virus | 168804177 | 2 |
| 97 | ORF94 | LbFV | 1148998795 | 2 |
| 98 | ORF96 | Lb | PQAT00000000 | 2 |
| 99 | ORF96 | Lh | RICB00000000 | 2 |
| 100 | ORF96 | Lc | JUFY01000000 | 2 |
| 101 | ORF96 | LbFV | 1148998797 | 2 |
| 102 | ORF107 | Lb | PQAT00000000 | 2 |
| 103 | ORF107 | Lh | RICB00000000 | 2 |
| 104 | ORF107 | Lc | JUFY01000000 | 2 |
| 105 | ORF107 | Glossina_pallidipes_salivary_gland_hypertrophy_virus | 168804057 | 2 |
| 106 | ORF107 | Musca_domestica_salivary_gland_hypertrophy_virus | 187903107 | 2 |
| 107 | ORF107 | LbFV | 1148998799 | 2 |

*Continued on next page*

| | Locus | species | GI | Figure |
|---|---|---|---|---|
| 108 | ITS2 | L.longipes | AF015893.1 | S19 |
| 109 | ITS2 | L.guineaensis | AY124559.1 | S19 |
| 110 | ITS2 | L.victoriae | AY124553.1 | S19 |
| 111 | ITS2 | L.heterotoma | AB546896.1 | S19 |
| 112 | ITS2 | L.orientalis | AY124563.1 | S19 |
| 113 | ITS2 | L.boulardi | AY124568.1 | S19 |
| 114 | ITS2 | L.freyae | AY124561.1 | S19 |
| 115 | ITS2 | L.fimbriata | AF015894.1 | S19 |
| 116 | ITS2 | L.clavipes | JQ808416.1 | S19 |
| 117 | ITS2 | L.australis | AF015897.1 | S19 |
| 118 | ITS2 | G.brasiliensis | AB678777.1 | S19 |
| 1 | ORF27 | Papilio xuthus | XP_013173302.1 | S1A |
| 2 | ORF27 | Bicyclus anynana | XP_023937808.1 | S1A |
| 3 | ORF27 | Pieris rapae | XP_022114989.1 | S1A |
| 4 | ORF27 | Spodoptera litura | XP_022828254.1 | S1A |
| 5 | ORF27 | Bombyx mori | NP_001037024.1 | S1A |
| 6 | ORF27 | Drosophila busckii | XP_017843635.1 | S1A |
| 7 | ORF27 | Musca domestica | XP_005178734.1 | S1A |
| 8 | ORF27 | Zeugodacus cucurbitae | XP_011180685.1 | S1A |
| 9 | ORF27 | Ceratitis capitata | XP_004519914.1 | S1A |
| 10 | ORF27 | Dendroctonus ponderosae | XP_019755885.1 | S1A |
| 11 | ORF27 | Anoplophora glabripennis | XP_018566786.1 | S1A |
| 12 | ORF27 | Leptinotarsa decemlineata | XP_023022306.1 | S1A |
| 13 | ORF27 | Polistes dominula | XP_015178412.1 | S1A |
| 14 | ORF27 | Linepithema humile | XP_012229104.1 | S1A |
| 15 | ORF27 | Camponotus floridanus | XP_011252805.1 | S1A |
| 16 | ORF27 | Pogonomyrmex barbatus | XP_011630441.1 | S1A |
| 17 | ORF27 | Megachile rotundata | XP_012151451.1 | S1A |
| 18 | ORF27 | Microplitis demolitor | XP_008554575.1 | S1A |
| 19 | ORF27 | Fopius arisanus | XP_011298329.1 | S1A |
| 20 | ORF27 | Diachasma alloeum | XP_015109162.1 | S1A |
| 21 | ORF27 | Cephus cinctus | XP_015599785.1 | S1A |
| 22 | ORF27 | Ganaspis brasiliensis | RJVV00000000 | S1A |
| 23 | ORF27 | Leptopilina boulardi | PQAT00000000.1 | S1A |
| 24 | ORF27 | Leptopilina heterotoma | RICB00000000 | S1A |
| 25 | ORF27 | Leptopilina clavipes | JUFY00000000.1 | S1A |
| 26 | ORF27 | Orussus abietinus | XP_012276925.1 | S1A |
| 27 | ORF27 | Nasonia vitripennis | XP_016838993.1 | S1A |
| 28 | ORF27 | LbFV | 1148998730 | S1A |
| 29 | ORF27 | Dufourea novaeangliae | XP_015432901.1 | S1A |
| 30 | ORF27 | Apis florea | XP_012348205.1 | S1A |
| 31 | ORF27 | Apis mellifera | XP_006570777.1 | S1A |
| 32 | ORF27 | Habropoda laboriosa | XP_017799036.1 | S1A |
| 33 | ORF27 | Bombus terrestris | XP_012163415.1 | S1A |
| 34 | ORF66 | Harpegnathos saltator | 749795708 | S1B |
| 35 | ORF66 | Camponotus floridanus | 752871224 | S1B |
| 36 | ORF66 | Pogonomyrmex barbatus | 769838565 | S1B |
| 37 | ORF66 | Vollenhovia emeryi | 795079157 | S1B |
| 38 | ORF66 | Nasonia vitripennis | 1032757220 | S1B |
| 39 | ORF66 | Trichomalopsis sarcophagae | 1227108847 | S1B |
| 40 | ORF66 | Cephus cinctus | 1000753753 | S1B |
| 41 | ORF66 | Ganaspis brasiliensis | RJVV00000000 | S1B |
| 42 | ORF66 | Leptopilina heterotoma | RICB00000000 | S1B |
| 43 | ORF66 | Leptopilina clavipes | JUFY00000000.1 | S1B |
| 44 | ORF66 | Leptopilina boulardi | PQAT00000000.1 | S1B |
| 45 | ORF66 | Dufourea novaeangliae | 987914045 | S1B |
| 46 | ORF66 | Habropoda laboriosa | 1059864473 | S1B |
| 47 | ORF66 | Apis florea | 820863019 | S1B |
| 48 | ORF66 | Apis dorsata | 572314547 | S1B |
| 49 | ORF66 | Apis mellifera | 571506210 | S1B |
| 50 | ORF66 | Apis cerana cerana | 1241837182 | S1B |
| 51 | ORF66 | Eufriesea mexicana | 1059214553 | S1B |
| 52 | ORF66 | Bombus terrestris | 240708910 | S1B |
| 53 | ORF66 | Agrilus planipennis | XP_018331076.1 | S1B |
| 54 | ORF66 | Tribolium castaneum | NP_001280519.1 | S1B |
| 55 | ORF66 | Nicrophorus vespilloides | XP_017784576.1 | S1B |
| 56 | ORF66 | Papilio machaon | 930680047 | S1B |
| 57 | ORF66 | Papilio xuthus | 910339325 | S1B |
| 58 | ORF66 | Helicoverpa armigera | 204423112 | S1B |
| 59 | ORF66 | Trichoplusia ni | 6635437 | S1B |
| 60 | ORF66 | Acyrthosiphon pisum | 228698707 | S1B |
| 61 | ORF66 | Diuraphis noxia | 985403395 | S1B |
| 62 | ORF66 | Myzus persicae | 1230193237 | S1B |
| 63 | ORF66 | LbFV | 1148998769 | S1B |
| 64 | ORF66 | Adoxophyes honmai EPV | 506498063 | S1B |
| 65 | ORF11-13 | LbFVorf11 | 009345615 | S1C |
| 66 | ORF11-13 | Ganaspis brasiliensis | RJVV00000000 | S1C |
| 67 | ORF11-13 | LbFVorf13 | 009345617.1 | S1C |
| 68 | ORF11-13 | Leptopilina boulardi | PQAT00000000.1 | S1C |
| 69 | ORF11-13 | Leptopilina heterotoma | RICB00000000 | S1C |
| 70 | ORF11-13 | Leptopilina clavipes | JUFY00000000.1 | S1C |
| 71 | ORF11-13 | Exserohilum turcica | XP_008030043.1 | S1C |
| 72 | ORF11-13 | Alternaria alternata | XP_018379425.1 | S1C |
| 73 | ORF11-13 | Frankliniella occidentalis | XP_026288761.1 | S1C |
| 74 | ORF11-13 | Rhizopus microsporus | XP_023462188.1 | S1C |
| 75 | ORF11-13 | Melampsora larici-populina | XP_007414376.1 | S1C |
| 76 | ORF11-13 | Debaryomyces hansenii | XP_459998.2 | S1C |
| 77 | ORF11-13 | Debaryomyces fabryi | XP_015465751.1 | S1C |
| 78 | ORF11-13 | Eremothecium gossypii | NP_986783.2 | S1C |
| 79 | ORF11-13 | Eremothecium cymbalariae | XP_003645815.1 | S1C |

| | primer_name | Orientation | tm | GC | Seq | Prod.Size |
|---|---|---|---|---|---|---|
| 1 | Lb_ORF96_F | FORWARD | 59.99 | 55 | AATGGAGGACTACCGACACG | 259 |
| 2 | Lb_ORF96_R | REVERSE | 59.62 | 47 | TGCACTGTGGTCCATAAACAG | |
| 3 | Lb_ORF92_F | FORWARD | 59.94 | 45 | TGACCAAGACATGGTGGAAA | 248 |
| 4 | Lb_ORF92_R | REVERSE | 60.07 | 45 | CCGAATTGAATGACATGCTG | |
| 5 | Lb_ORF58_F | FORWARD | 59.65 | 50 | TACCAAATGGTGGAGGGAAC | 250 |
| 6 | Lb_ORF58R | REVERSE | 59.60 | 40 | CCATTTAAAACGTCGCAACA | |
| 7 | Lb_ORF68F | FORWARD | 59.79 | 50 | TGTCTGGAGATTGCCATCAG | 239 |
| 8 | Lb_ORF68R | REVERSE | 60.04 | 45 | CCAATTTTCGGAAGTGAGGA | |
| 9 | Lb_ORF5F | FORWARD | 60.41 | 40 | GATTCGCCAAATTTGATTGC | 243 |
| 10 | Lb_ORF5R | REVERSE | 60.08 | 45 | ATCATCATTGTCAGCGTCCA | |
| 11 | Lb_ORF60F | FORWARD | 59.89 | 50 | ACGTACGATTGGCGTAAACC | 235 |
| 12 | Lb_ORF60R | REVERSE | 60.84 | 55 | GACGTTGTTGTCCGAAGAGC | |
| 13 | Lb_ORF85F | FORWARD | 59.77 | 40 | CAGCTTTAGAACCTTGGGAAAA | 249 |
| 14 | Lb_ORF85R | REVERSE | 59.73 | 45 | GCCAACGCTGCACATTATTA | |
| 15 | Lb_ORF78F | FORWARD | 60.07 | 45 | CGATTTTGATGGTGATGCAG | 251 |
| 16 | Lb_ORF78R | REVERSE | 59.31 | 40 | CATTTTCAATGCACGAAAGC | |
| 17 | Lb_ORF94F | FORWARD | 60.22 | 45 | TGCCGTCGAAGATACATTCA | 252 |
| 18 | Lb_ORF94R | REVERSE | 58.85 | 50 | TCCACGGTAGACCATGTGTT | |
| 19 | Lb_ORF107F | FORWARD | 59.62 | 55 | CGACGCTATTGCAGTCAGTC | 251 |
| 20 | Lb_ORF107R | REVERSE | 60.00 | 45 | GCGTCAGAAGCAACAAATGA | |
| 21 | Lb_ORF87F | FORWARD | 60.21 | 35 | TTGCAATATGCCAGCAAAAA | 260 |
| 22 | Lb_ORF87R | REVERSE | 59.92 | 40 | GTTCCCAGGCAAAAATTTCA | |
| 23 | Lb_ORF72F | FORWARD | 59.96 | 45 | CTTTTTGCGGATCTTTCAGC | 236 |
| 24 | Lb_ORF72R | REVERSE | 60.66 | 55 | CTCCATTCTTGCCTGGACAC | |
| 25 | Lb_ORF83F | FORWARD | 56.00 | 40 | ATTCCAATGGTTGGCGAATA | 84 |
| 26 | Lb_ORF83R | REVERSE | 62.00 | 55 | CCGAGTGGAGTACACGTTTG | |
| 27 | Lb_RhoGapF | FORWARD | 56.00 | 40 | AATTCGGAAGCAATGGAAGA | 325 |
| 28 | Lb_RhoGapR | REVERSE | 56.00 | 40 | ATCGCTTGGTTTCTTTTTGC | |
| 29 | Lb_actinF | FORWARD | 66.00 | 65 | GATGCCCCGAGGCTCTCTTC | 294 |
| 30 | Lb_actinR | REVERSE | 60.00 | 52 | TGGTGCCAAGGCAGTGATT | |
| 31 | Lb_shakeF | FORWARD | 64.00 | 60 | CGAGTTATCGGTGCGCTTCC | 182 |
| 32 | Lb_shakeR | REVERSE | 62.00 | 55 | GCGAGGGACATCGCTTGATT | |

Table S6: Primers used in the paper.

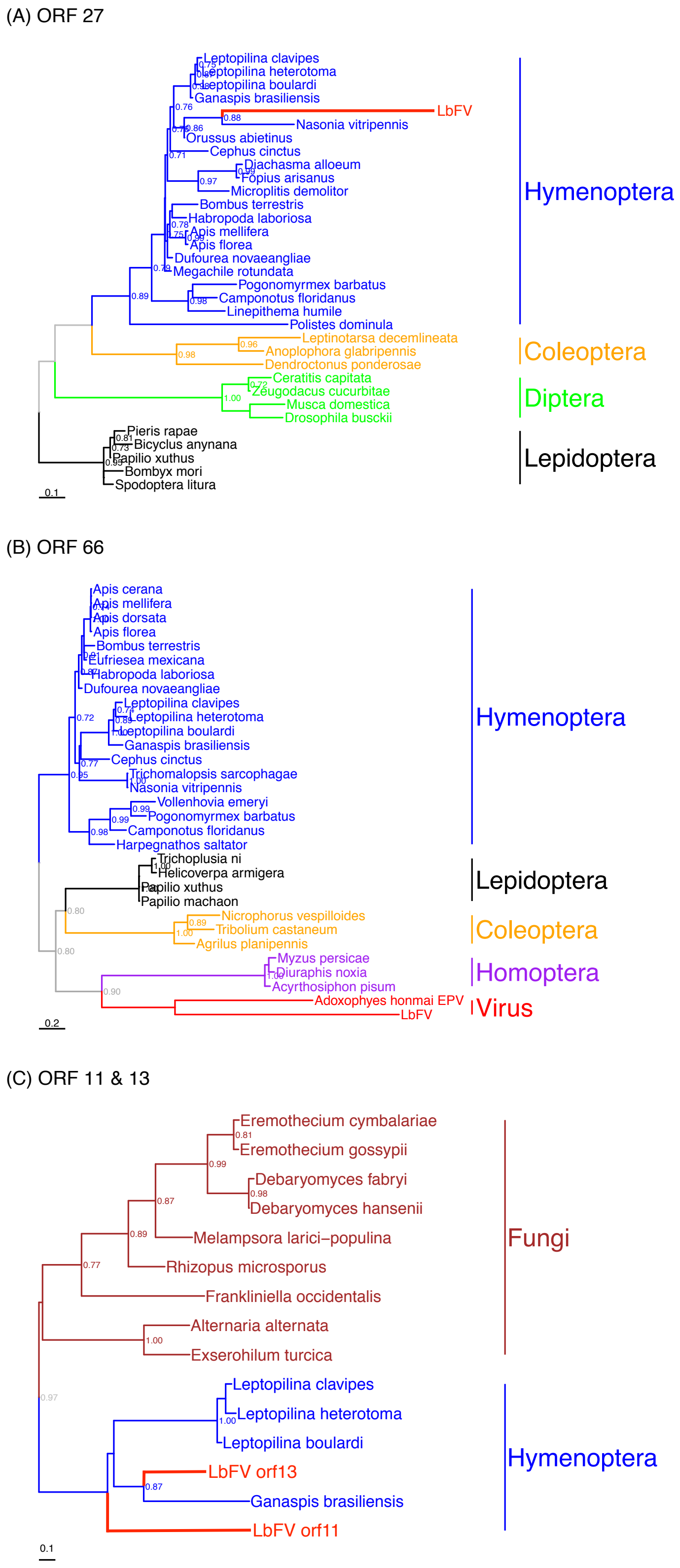


Figure S1: Four loci in the LbFV genome probably derive from insect genes. ORFs 27 (A) and 66 (B) are putative inhibitors of apoptosis and ORF 11 and 13 (C) contain a putative histone demethylase domain [76]. Sequences were aligned using muscle, and conserved blocks were identified using gblocks to construct a PhyML phylogeny (parameters: -d aa -m LG -b -4 -v e -c 4 -a e -f m). Only aLRT values ⩾ 0.7 are shown. Accession numbers of the corresponding sequences are available in table S5.

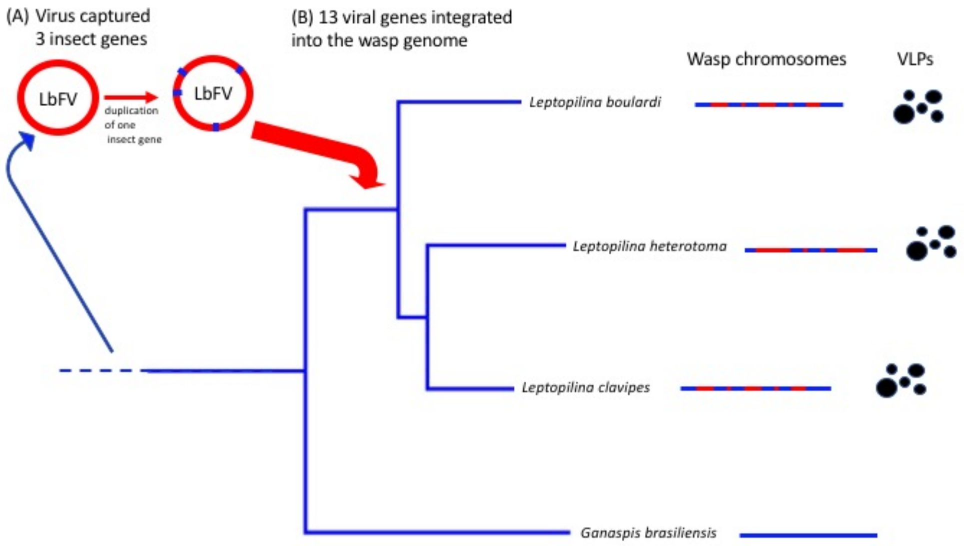


Figure S2: Hypothetical scenario for genetic exchanges between the wasps and the virus LbFV. (A) Before the diversification of the *Leptopilina* genus, LbFV captured 3 insect genes, most likely involved in apoptosis inhibition (ORFs 27 and 66) and methylation (the ancestor of ORFs 11 and 13). One of them was probably subsequently duplicated (the ancestor of both ORFs 11 and 13). (B) After the divergence between *Ganaspis* and *Leptopilina* (around 74My ago[9]), but before the diversification of *Leptopilina* genus, possibly a whole genome of a virus closely related to LbFV integrated wasp chromosomes. Nowadays, all *Leptopilina* species bear 13 LbFV-derived genes that allow them to produce VLPs. The cartoons displaying the chromomosomes are just illustrations depicting the presence of virally-derived genes (red) within wasp chromosomes of eukaryotic origin (blue). VLPs are symbolized by the black circular forms.

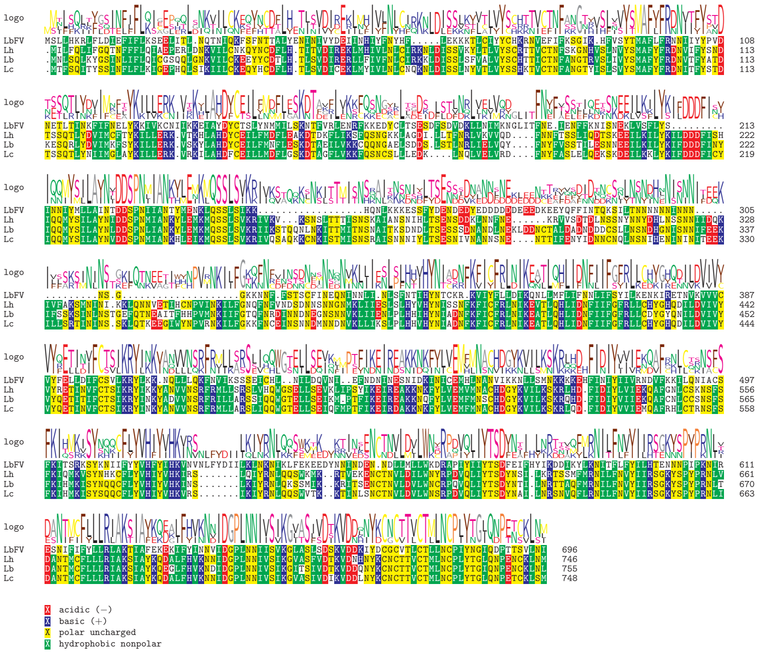


Figure S3: Alignment of LbFV ORF5 and their homologs in *Leptopilina*. Plot obtained using the msa R package[6].

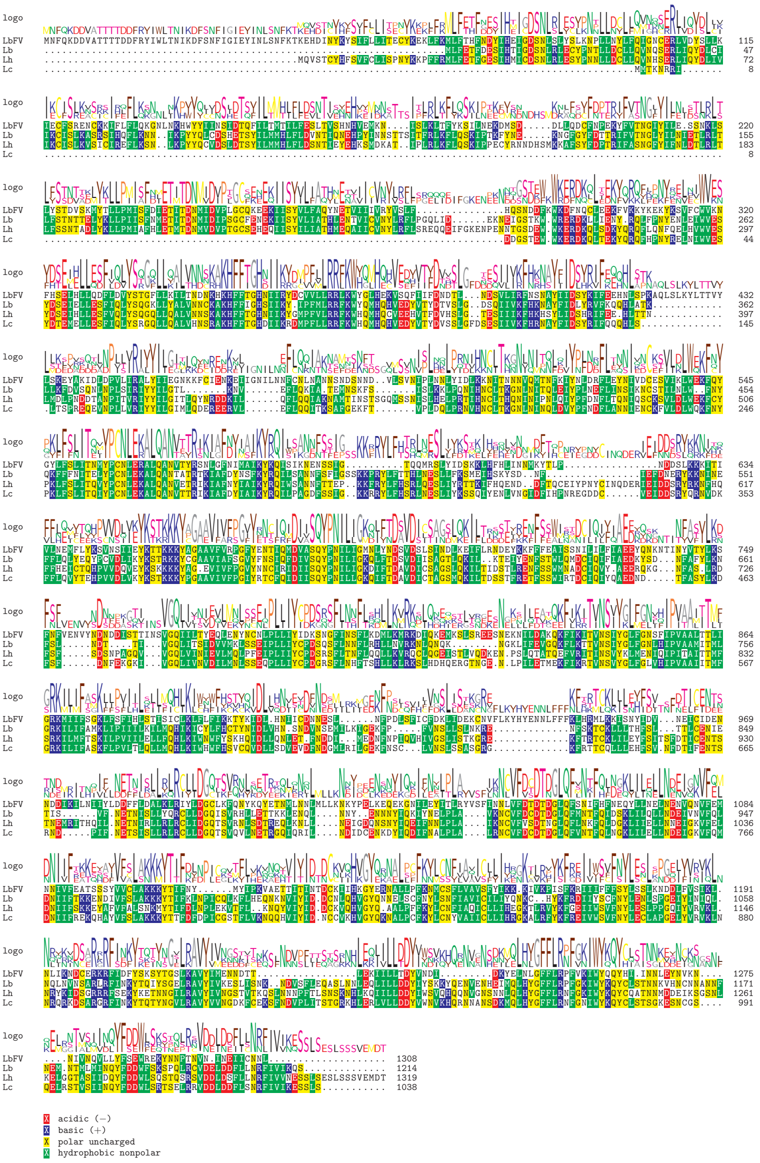


Figure S4: Alignment of LbFV ORF58 and their homologs in *Leptopilina*.

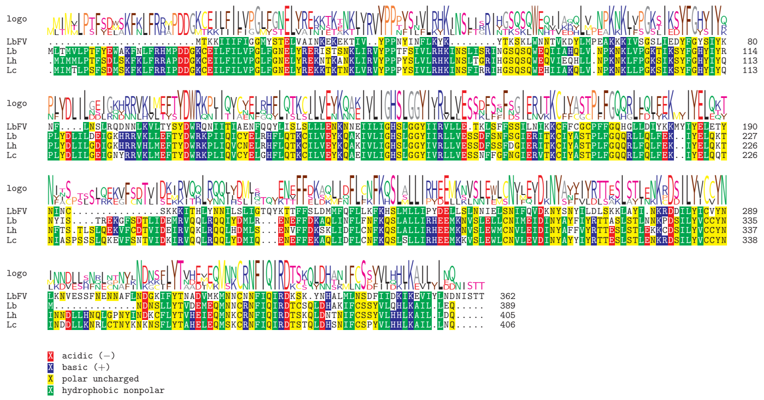

Figure S5: Alignment of LbFV ORF60 and their homologs in *Leptopilina*.

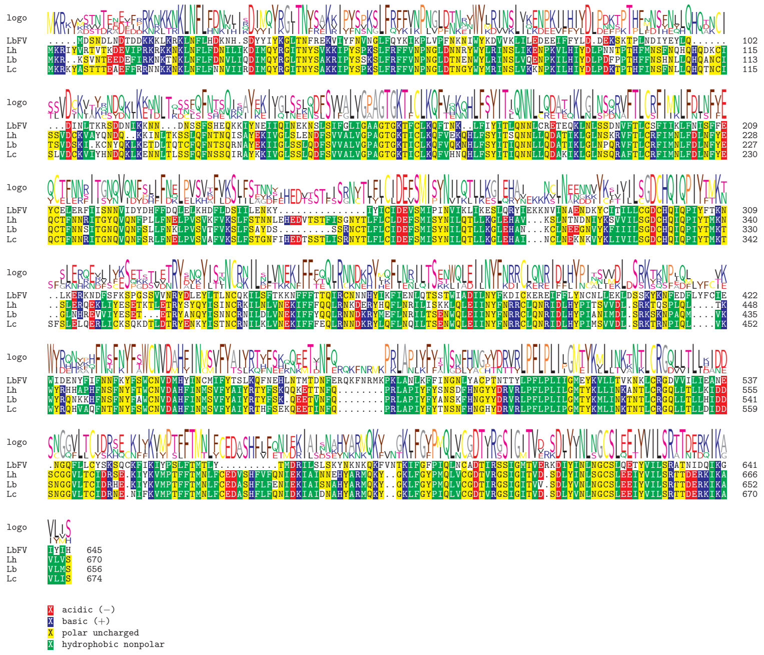


Figure S6: Alignment of LbFV ORF68 and their homologs in *Leptopilina*.

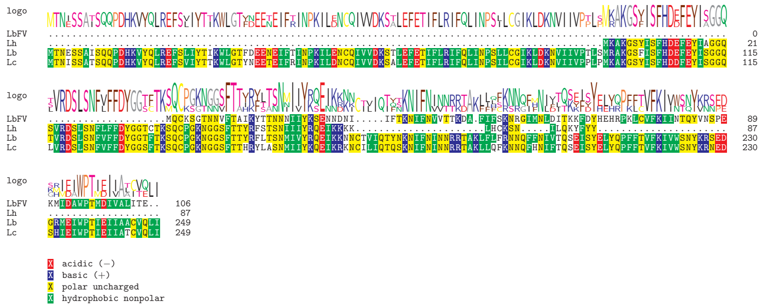

Figure S7: Alignment of LbFV ORF72 and their homologs in *Leptopilina*.

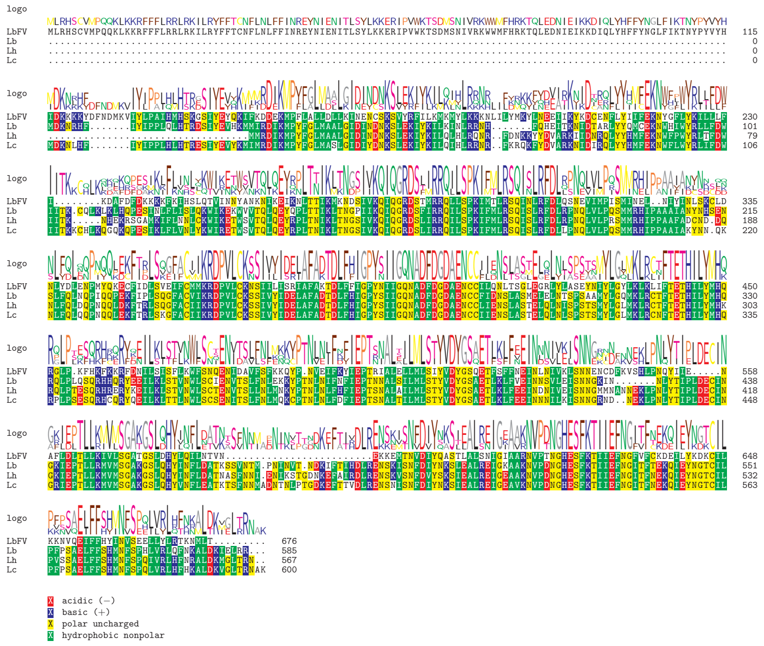

Figure S8: Alignment of LbFV ORF78 and their homologs in *Leptopilina*.

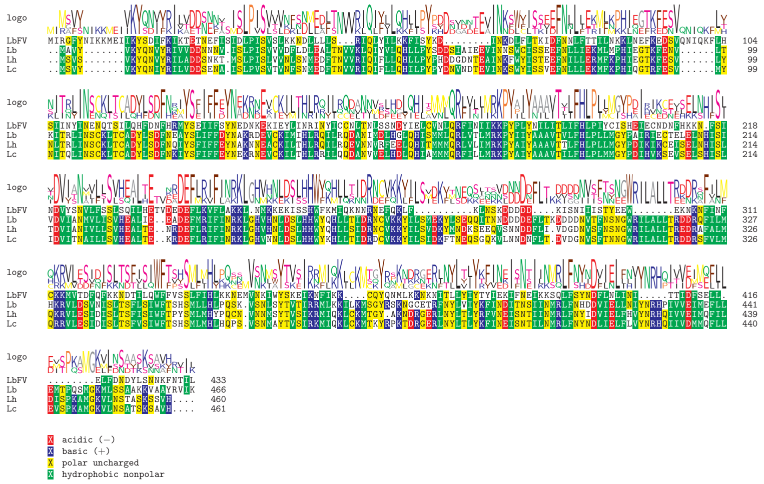


Figure S9: Alignment of LbFV ORF83 and their homologs in *Leptopilina*.

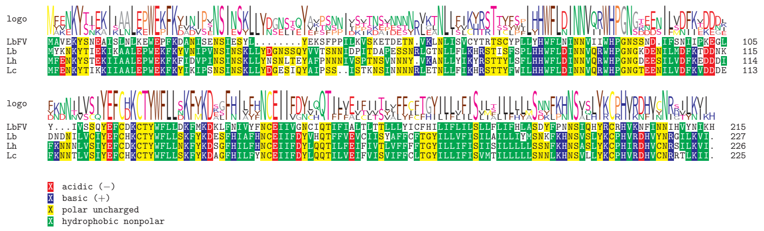

Figure S10: Alignment of LbFV ORF85 and their homologs in *Leptopilina*.

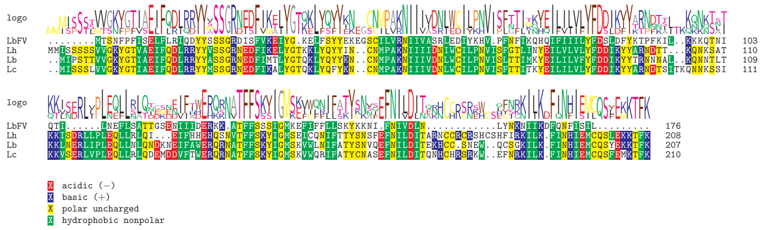

Figure S11: Alignment of LbFV ORF87 and their homologs in *Leptopilina*.

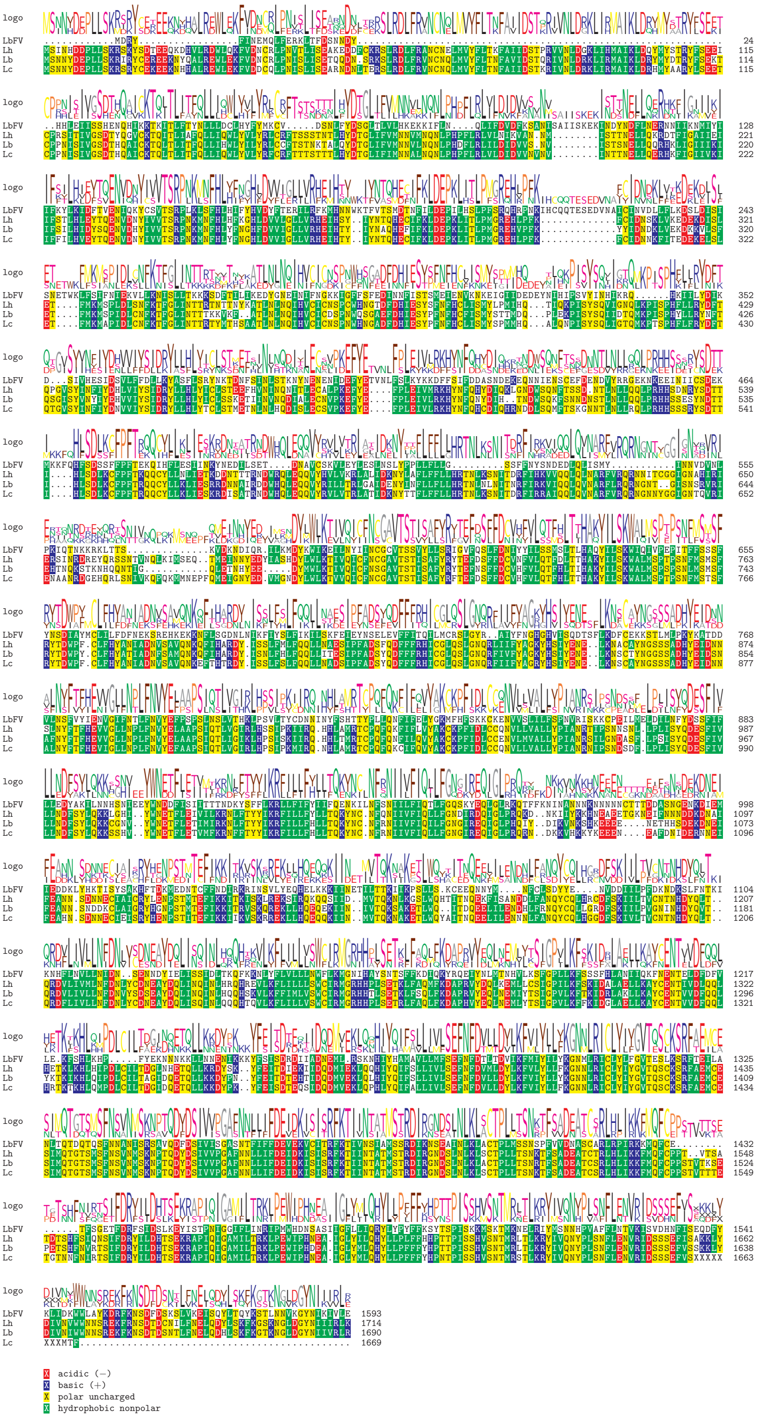

Figure S12: Alignment of LbFV ORF92 and their homologs in *Leptopilina*.

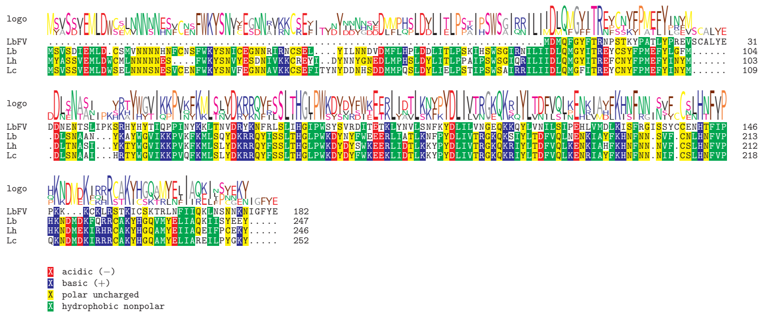


Figure S13: Alignment of LbFV ORF94 and their homologs in *Leptopilina*.

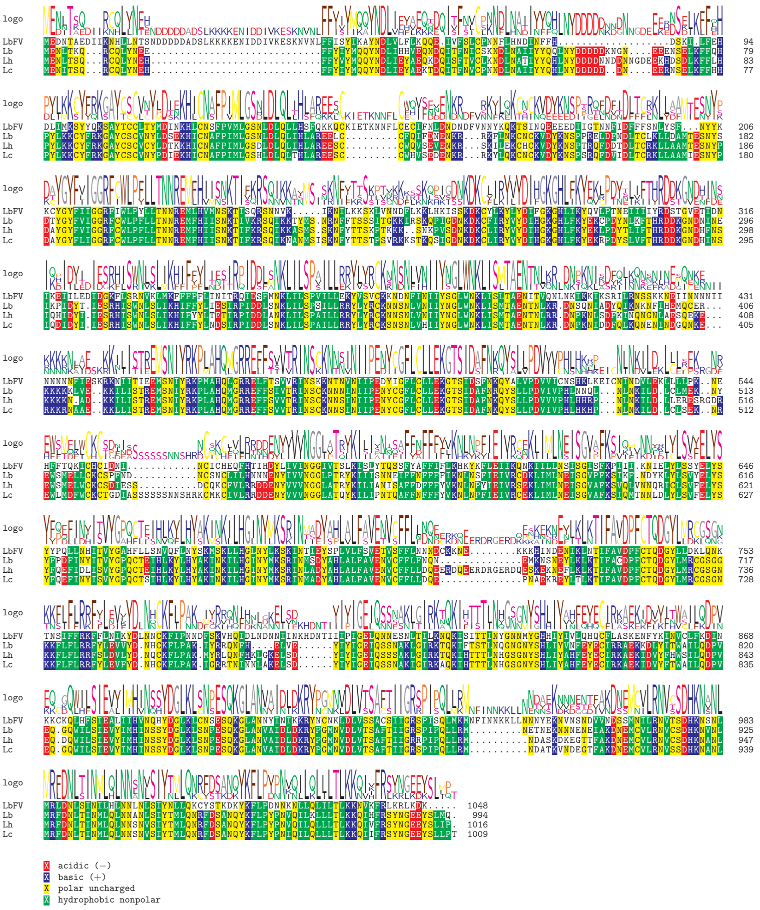


Figure S14: Alignment of LbFV ORF96 and their homologs in *Leptopilina*.

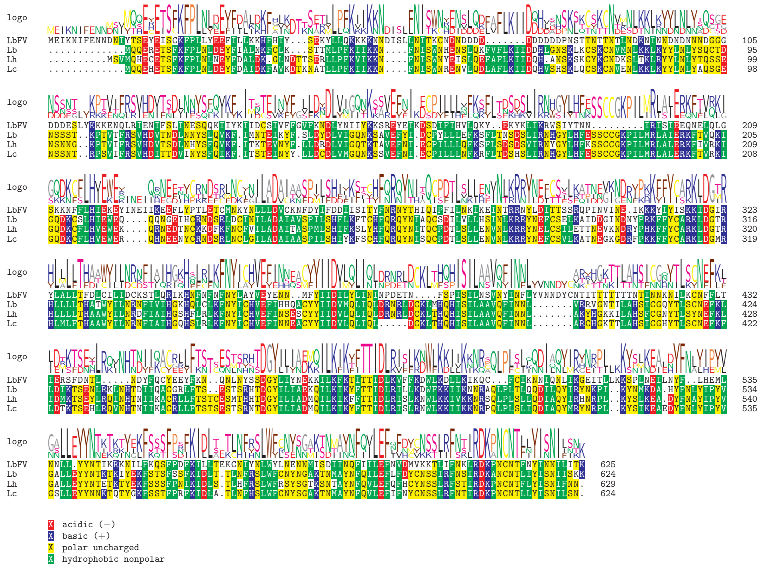

Figure S15: Alignment of LbFV ORF107 and their homologs in *Leptopilina*.

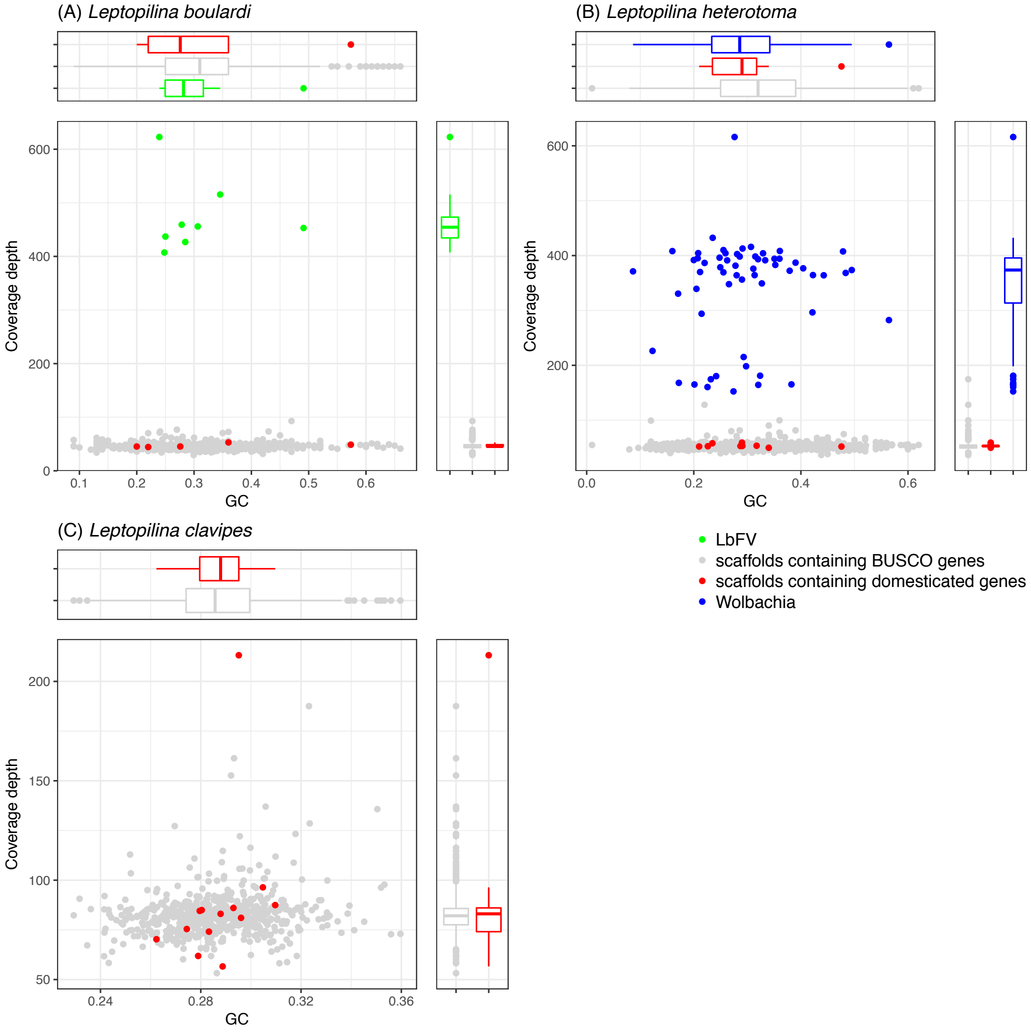


Figure S16: General features of scaffolds containing single copy universal arthropod genes (BUSCO gene set, in grey), scaffolds containing virally-derived loci (in red), scaffolds belonging to the virus LbFV (in green, only in *L. boulardi*) and of scaffolds belonging to *Wolbachia* endosymbiont (in blue, only in *L. heterotoma*). The heterogeneity in coverage depth for the *Wolbachia* scaffolds in *L. heterotoma* is probably the consequence of multi-infection with three *Wolbachia* strains having different densities[59]. (A) *L. boulardi*; (B) *L. heterotoma*, (C) *L. clavipes*.

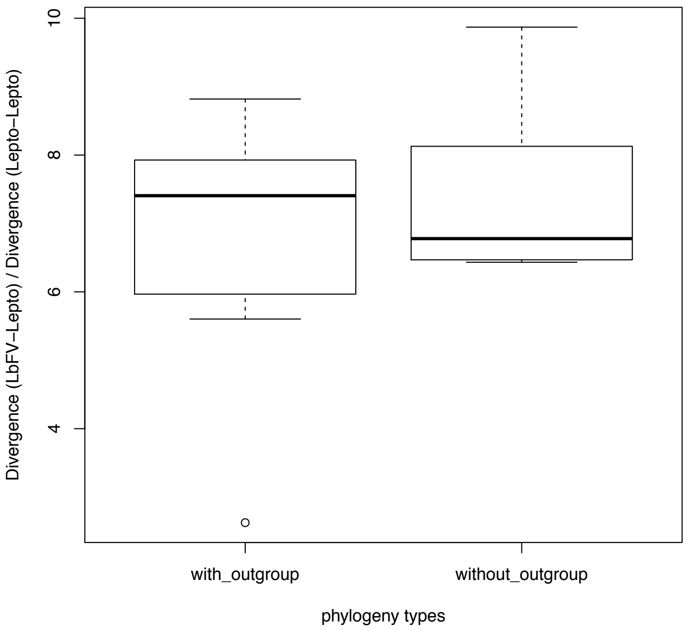


Figure S17: Divergence of LbFV with *Leptopilina* species relative to the divergence among *Leptopilina* species. This relative divergence was calculated both for the seven loci for which additional viral sequences were found (in addition to the LbFV sequence, "with_outgroup") and for the six loci for which no additional viral sequences were found ("without_outgroup"). The relative divergence is not statistically different between phylogeny types (F(1,1)=0.9, p-value=0.37). This further suggests that the all 13 genes have the same evolutionary history.

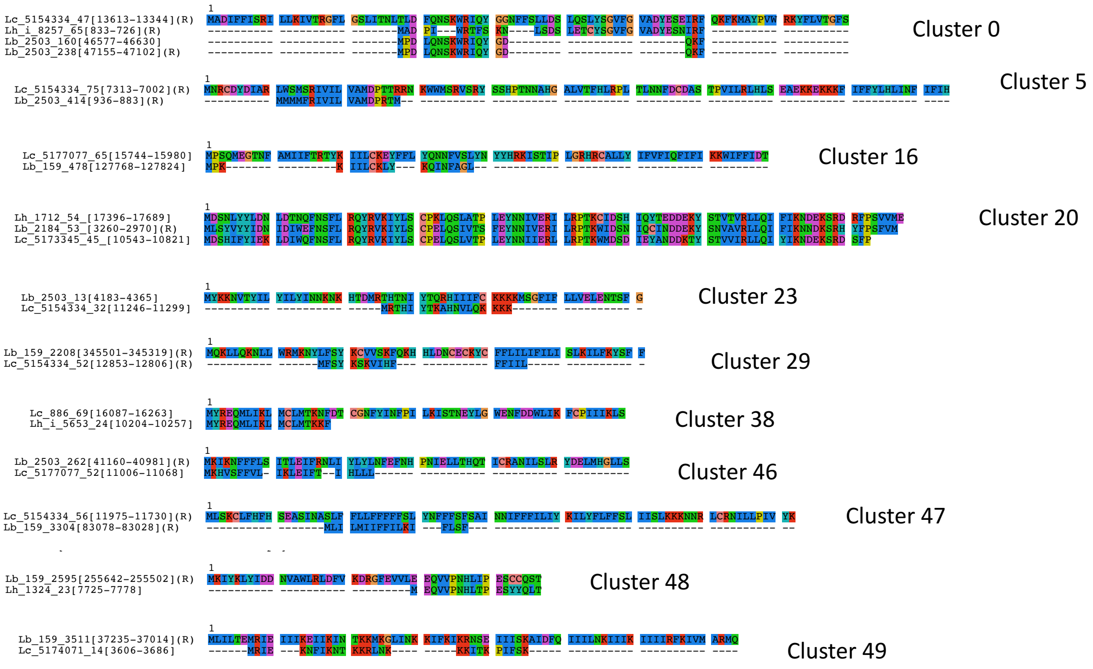


Figure S18: Flanking regions of virally-derived genes show similarities between *Leptopilina* species. Amino-acid sequences were predicted from the wasps scaffolds containing the virally-derived genes (but masked for the viral genes themselves) using getorf (-minsize 50 -find 1). They were clustered using CD-hit (-c 0.7), and aligned using muscle.

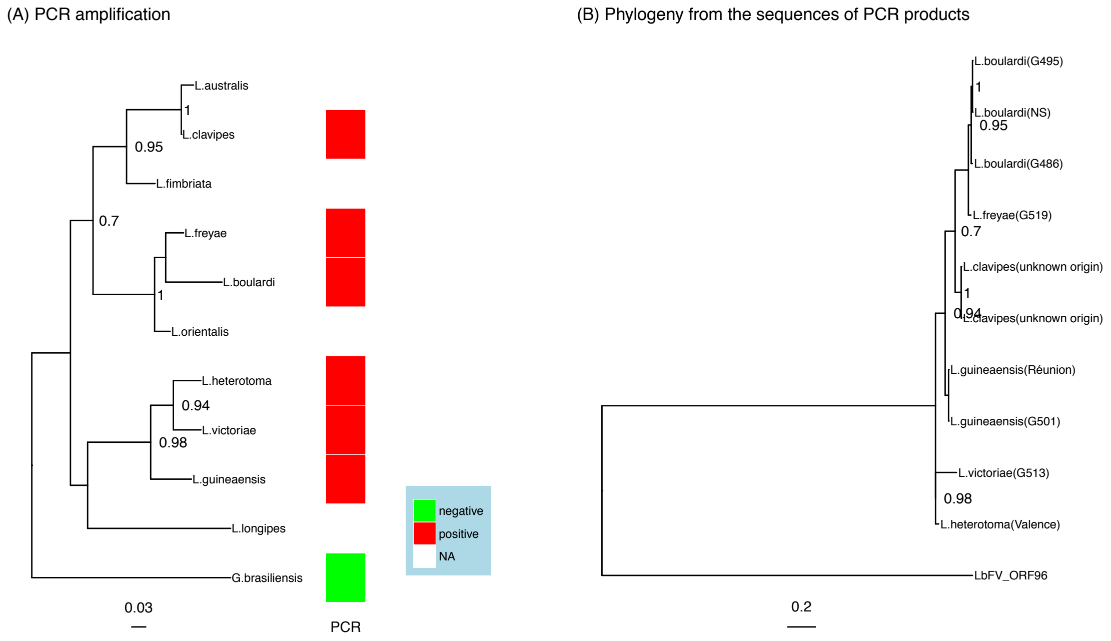


Figure S19: Amplification, sequencing and phylogeny of orthologs of LbFV_ORF96 in *Leptopilina* species. (A) Phylogeny of *Leptopilina* genus and *Ganaspis brasiliensis* based on internal transcribed spacer 2 (ITS2). (B) Phylogeny obtained after sequencing the corresponding PCR products. The strain used is indicated between brackets. Only aLRT $\geqslant 0.70$ are shown. Accession numbers of the corresponding sequences are available in table S5.

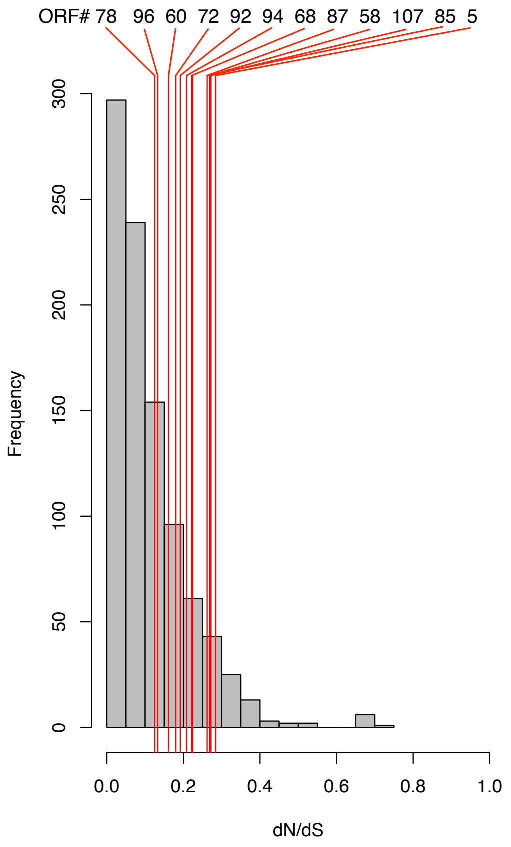


Figure S20: dN/dS ratio for a set of 942 universal arthropod genes and for the 13 virally derived genes found in *Leptopilina* species (indicated by the red lines).

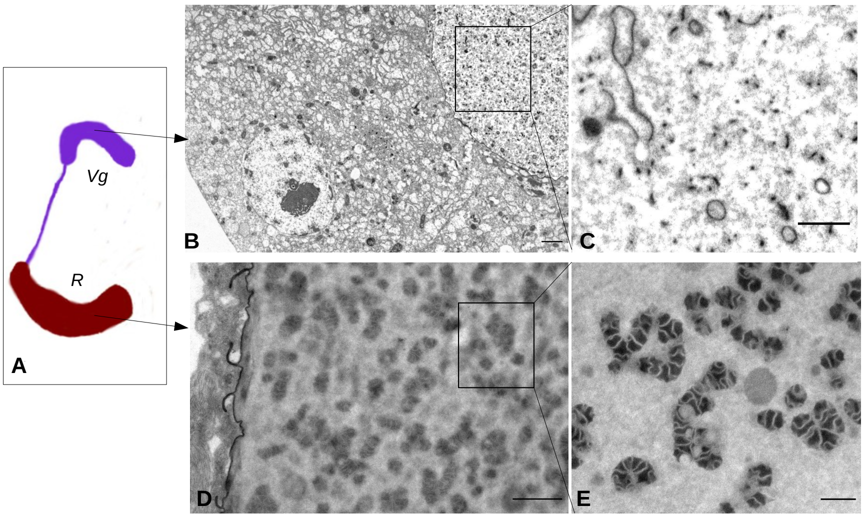


Figure S21: Similarly to other *Leptopilina* studied so far, VLPs are produced in the venom gland of *L. clavipes.* A venom gland cartoon is displayed in (A) with the venom gland itself (Vg) where VLPs are produced and the reservoir (R) where mature VLPs acumulate. A cross section of the venom gland observed under transmission electron microscopy reveals the presence of membranes in the lumen (B-C). Their acquire the typical structure of mature VLPs in the reservoir (D-E). Bars represent : 2 $\mu$M (B); 1 $\mu$M (C); 1 $\mu$M (D) and 0.5 $\mu$M (E) 1$\mu$M.

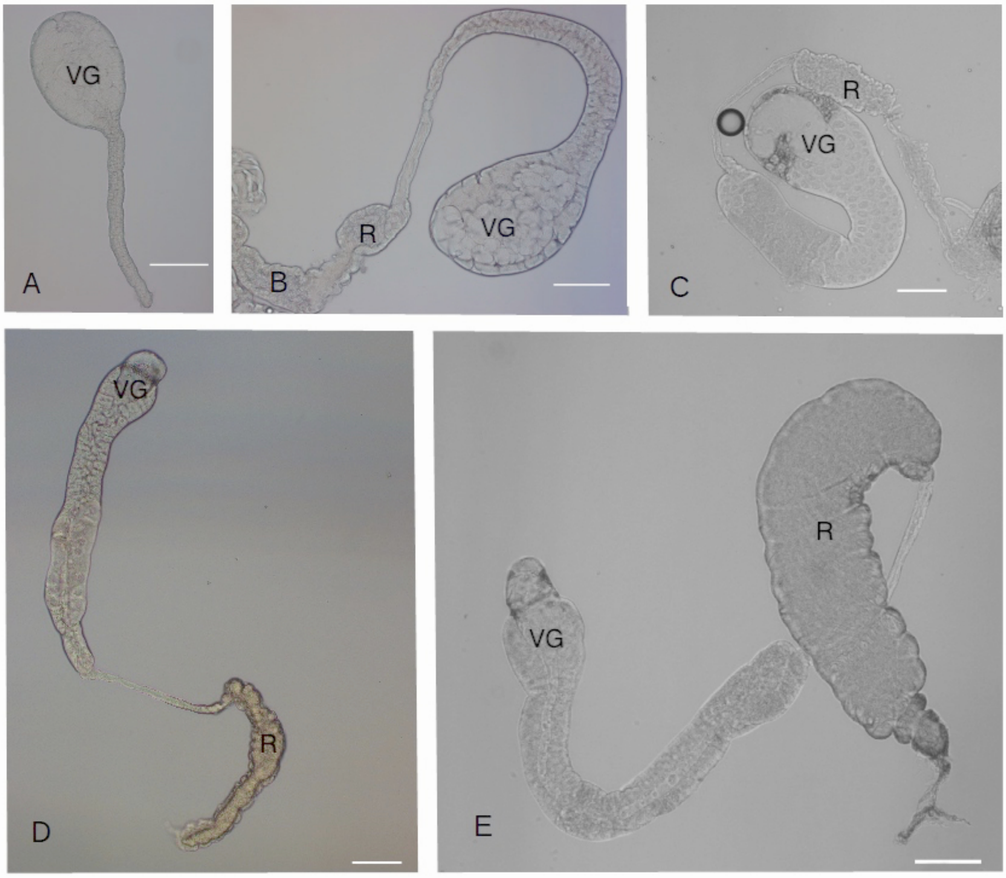


Figure S22: Morphogenesis of the venom gland during the pupal stage of *L. boulardi* females. G: venom gland; R: reservoir of the venom gland. Overall structure of the organ under light microscope at day 11 (a), 14 (b), 16 (c), 18(d) and 21(e). At that temperature (25°C), 11 days corresponds to the beginning of the pupation in *L. boulardi*, whereas adult females are emerging at 21 days. Bar= 100$\mu$M.

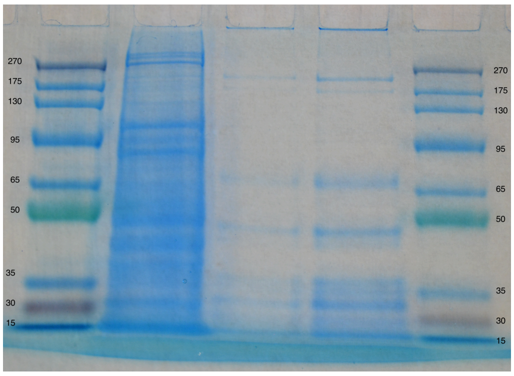


Figure S23: SDS-PAGE gel of proteins from purified VLPs. Lane 1 : ladder, Lane 2: positive control (whole drosophila), lane 3: VLP sample 1, lane 4: VLP sample 2, lane 5: ladder.

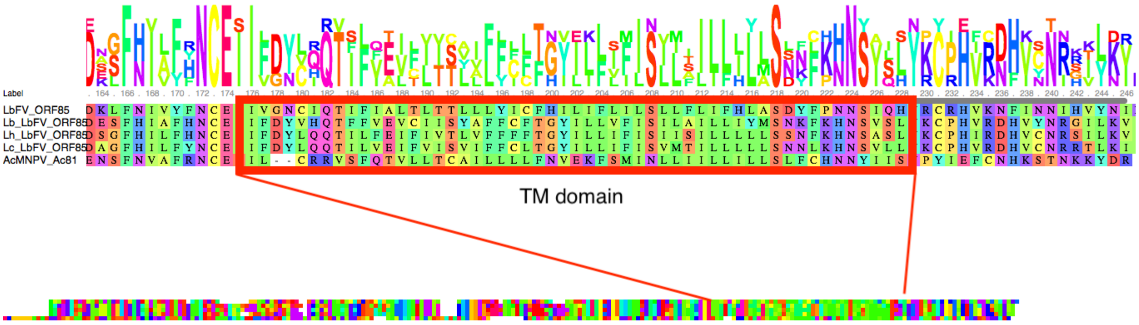


Figure S24: Ac81 homologs in LbFV and in *Leptopilina* genomes (ORF85) share a conserved hydrophobic, probably transmembrane domain.
